# The reproduction process of Gram-positive protocells

**DOI:** 10.1101/2021.11.25.470039

**Authors:** Dheeraj Kanaparthi, Marko Lampe, Jan-Hagen Krohn, Baoli Zhu, Falk Hildebrand, Thomas Boesen, Andreas Klingl, Prasad Phapale, Tillmann Lueders

## Abstract

Prokaryotes are hypothesized to have evolved from more primitive protocells. Unlike present-day cells, protocells are thought to have been devoid of complex molecular biological processes. They are believed to have mediated reproduction entirely by biophysical forces under favorable environmental conditions. Despite this proposition, little is known about the actual mechanism of their reproduction. To understand the reproduction process of protocells in their native habitat, here we used a top-down approach to transform bacterial cells into a primitive lipid vesicle-like state. Given that environmental conditions are thought to have played an essential role in mediating protocell reproduction, we then studied these cells under the presumed environmental conditions of Archaean Eon Earth. Even in the absence of functioning biological processes, cells in our study reproduced in a defined sequence of steps, always leading to the formation of viable daughter cells. Their reproduction mechanism can be explained by the interaction between intracellular metabolism, physicochemical properties of cell constituents, and, most importantly, environmental conditions. Given the simplicity of this reproduction mechanism and its suitability to environmental conditions of early Earth, we propose that protocells reproduced by this process. Moreover, this method of reproduction is also in tune with the earlier theoretical propositions on protocells, the results of the top-down approach of building a minimal cell, and the paleontological record of the Achaean Eon. Our study is the first to bridge the gap between non-living systems like lipid vesicles, living cells, and the paleontology of the Archaean Eon.

## Introduction

The theory of chemical evolution proposes that prokaryotes evolved from more primitive protocells (1). Molecular biological processes in these primitive cells are hypothesized to have been simpler and far less complex than the present-day cells (2, 3). Because protocells lack complex molecular biological processes, they were believed to reproduce through simple biophysical processes under favorable environmental conditions (3–5). However, much of what we know about protocells has been derived from studying lipid vesicles (used as model protocells) under well-controlled laboratory settings (4, 5). Despite achieving significant progress, we believe lipid vesicles (LVs) do not represent the true physiological complexity of protocells. Moreover, studying an organism outside its natural habitat also fails to provide a complete picture of its ecological function and behavior (6). This is especially true for protocells, given their greater dependence on environmental conditions to mediate essential cellular functions like reproduction. To understand the reproduction of protocells in their natural habitat, here we worked with protoplasts of Gram-positive bacteria as model protocells (*Exiguobacterium strain M* protoplasts *–EM-P*) under the environmental conditions of early Earth.

Our choice of using bacterial protoplasts is based on three key considerations. First, protocells like bacterial protoplasts were likely devoid of a peptidoglycan cell wall. Synthesis of individual cell wall components, transporting them across the cell membrane, and assembling them into a cell wall is a physiologically complex process requiring many enzymes working in a coordinated manner (7, 8). Evolutionarily primitive protocells most certainly are considered to have been devoid of such complex intracellular processes (4).

Moreover, high-resolution FIB-SEM imaging of the earliest known microfossils was shown to be surrounded by a 5nm carbon layer (9) resembling the thickness of a cell membrane rather than a much thicker cell wall(10). Second, like the theoretical propositions on protocells, protoplasts lack an innate ability to regulate their morphology or mediate reproduction (11–13). They can be described as a sack of cytoplasm reproducing in a haphazard manner (14). Third, even the most primitive cell must possess three essential characteristics – biochemical processes for harvesting free energy, a mechanism of using this energy to sustain minimal biosynthetic processes, and a mechanism of reproduction involving the transfer of its genetic material to the daughter cells. Even these basic processes incorporates considerable cytoplasmic complexity. Therefore, the biophysical properties of protocells may be more similar to bacterial cytoplasm than compositionally simpler LVs.

Given these similarities between protocells and protoplasts, we argue that bacterial protoplasts could be a better proxy than LVs for studying protocell reproduction.

Self-assembly of lipids into LVs happens within a narrow pH range and is influenced by environmental conditions like temperature and salt content (15). Hence, most protocell studies are conducted under narrow range of environmental conditions. Despite the theoretical propositions on protocells suggesting that their reproduction is greatly dependent on the environmental conditions, currently used experimental conditions rarely resemble the natural habitat of protocells. To understand the protocell reproduction in its native habitat, we grew *EM-P* under environmental conditions mimicking those that could have been prevalent on early Earth. The exact environmental conditions of Archaean Earth are currently unclear. Nevertheless, an emerging consensus among the scientific community suggests that at least by 3.8 billion years ago, surface temperatures likely ranged between 26° and 35°C (16, 17).

Early Earth’s oceans are also hypothesized to be 1.5 to 2 times saltier than the present-day oceans (18). Moreover, all known paleontological evidence of life is restricted to Archaean Eon coastal marine environments (19). This suggests primitive life (if not the very first cells) thrived under saline conditions. To replicate these conditions, we grew *EM-P* in half-strength tryptic soy broth with 7% (w/v) Dead Sea Salt (TSB-DSS) at 30°C. We used DSS, rather than pure NaCl, in our experiments to better simulate the complex salt compositions of natural environments. We then monitored these cells at regular intervals using a variety of super-resolution microscopic techniques to understand if and how these cells grow and reproduce. In the following sections of the manuscript, we present the lifecycle of *EM-P* under Archaean Eon environmental conditions, followed by a mechanistic explanation.

## Methods

*Exiguobacterium* strain-Molly (*EM*) was isolated from submerged biofilms in the Dead Sea (20). Protoplasts of *Exiguobacterium* Strain-Molly (*EM-P*) were generated by growing the cells in 7%DSS-TSB with Lysozyme and Penicillin G (Sigma Aldrich, Germany). Transformation to protoplast was confirmed by a change in cell morphology (Carl Zeiss, Germany). Given the stability of protoplasts, the use of both lysozyme and penicillin G was discontinued for subsequent experiments. For some experiments, *EM-P* was also grown in TSB with 4%DSS, 15%DSS, 6%MgCl_2_, and 7%KCl (Sigma Aldrich, Germany). We also grew the cells in 3X TSB containing 0.3M sucrose, 0.5% glycerol, 300mg of BSA, and 20mg of extracellular DNA.

Physiological characterization of *EM* was done in aerobic and anaerobic minimal salt media (21) with 2% w/v glucose as a sole carbon source. Aerobic and anaerobic media were inoculated (500ml) with overnight cultures of *EM*. A similar media composition was used to cultivate *EM-P* under aerobic and anaerobic conditions, except for adding 7%DSS. All the bottles were incubated on an orbital shaker at 30^0^C. All experiments were done in replicates (n=5). OD was determined every 3h interval for 15h. 1ml of the culture was centrifuged, and the supernatant was filter sterilized and stored at -20C for volatile fatty acid analysis (22).

Glucose was quantified using a Glucose colorimetric assay kit (Invitrogen, Germany) according to manufacturers’ instructions. Catalase activity was determined using the 19160 SOD detection kit (Sigma Aldrich, Germany). For anaerobic cultures, OD of all the test cultures was adjusted to similar values by dilution in original media, and cells were lysed by bead beating within the anaerobic chamber (Braun, Germany). Cell lysates were then taken out to measure catalase activity. All the experiments were done in replicates n=10 for the data presented in the manuscript, but the experiment was repeated several times throughout our work.

TEM and SEM were carried out using a Zeiss EM 912 (Carl Zeiss, Germany) with an integrated OMEGA filter as described before (23). Fluorescence microscopy was performed using an Olympus SpinSR10 spinning disk confocal microscope (Olympus, Tokyo, Japan). STED super-resolution fluorescence imaging was performed with an inverted TCS SP8 STED 3X microscope (Leica Microsystems, Germany) using 86x/1.2 NA water immersion objective (Leica HC PL APO CS2 - STED White).

To understand the relative contribution of intracellular processes and environmental conditions on the *EM-P*’s surface instability, we treated cells either with 2% methanol-free formaldehyde of 0.5% glutaraldehyde (ThermoFisher Scientific, Germany) to polymerize and immobilize the movement of proteins inside the cell. To equilibrate the conditions in and out of the cells, cells were permeabilized using 0.3% Triton X-100 (Sigma-Aldrich, Germany) for 10 min.

To quantify extracellular and intracellular DNA, we inoculated *EM-P* into the respective media. We incubated them either under static conditions or on an orbital shaker at 160 rpm. 2 ml of cells were harvested every day (24h) from all the incubations. Intact cells were precipitated by centrifugation at 6000 rpm for 10 min. DNA was extracted from both cells and supernatant using phenol-chloroform extraction. The resultant DNA was dissolved in 0.2 ml of TE buffer. Quantification of DNA was done using the Qubit dsDNA quantification kit (Invitrogen, Germany).

The viability of daughter cells was determined by inoculating 10-20 day-old cultures into fresh media either directly or after passing them through a 0.45μm cellulose acetate filter (Schematic shown in Figure 8) to get rid of large cells and membrane debris. Cytoplasmic activity in daughter cells was tested by staining them with DNA stain, PicoGreen^TM,^ and live cell labeling CellTrace^TM^ violet stain (Invitrogen, Germany) and imaging the cells using a confocal microscope. The relative percentage of cells with and without cytoplasmic activity and intracellular DNA was quantified by Attune CytPix flow cytometer (Thermo Fischer Scientific, Germany). The metabolic viability of *EM-P* daughter cells was also tested using the minimal salt media described above, with glucose as the sole carbon source.

## Results and Discussion

Given the lack of a cell wall, protoplasts were known to exhibit random morphologies and reproduce in a chaotic manner (11, 14). This reproduction method is shown to have been independent of canonical molecular biological processes that govern bacterial reproduction (11). In addition to losing control over its morphology and reproduction, more recent studies demonstrated that the loss of cell wall has more profound implications on cell physiology. These studies reported several discrepancies in intracellular functions, like an uncontrolled synthesis of cell wall precursors, even among stable protoplasts (24), the inability of protoplasts to maintain a proper intracellular oxidative state (25), uncontrolled replication of its genome (26) and excessive lipid (12). These studies suggest that cells in their protoplast state lack proper coordination between different cellular processes.

*EM-P* exhibited similar behavior. Cells in our incubations were in a perpetual state of lipid (membrane) excess (Fig. S1). The presence of excess membrane, together with the limited cytoplasmic volume of the cell, resulted in the formation of hollow extracellular and intracellular membrane structures as a mechanism of accommodating this excess membrane. We often observed log-phase *EM-P* cells with excess lipids within the cytoplasm in the form of lipid globules (Fig. S1a&b). Together with excessive lipid synthesis, *EM-P* also exhibited reduced efficiency in using terminal electron acceptors (Fig. S2) and a chaotic catalase expression (Supplementary results and discussion) compared to the wild-type cells with a cell wall. In tune with previous studies, these results suggest a lack of coordination among the processes in *EM-P*. However, in sharp contrast to the earlier studies (14), this loss of coordination did not result in an entirely random reproduction method. Under the environmental conditions of early Earth, *EM-P* reproduced in a defined sequence of steps, always leading to the formation of viable daughter cells. This consistency in its behavior was not due to the influence of molecular biological processes within the cell. Instead, it could be explained by the biophysical properties of its cell constituents, like the growth rate, excess lipid synthesis, and, most importantly, the environmental conditions.

Over the course of its life cycle, we observed *EM-P* reproducing by different processes. During the lag and early-log phase, cells remained spherical and gradually grew by an order of magnitude in diameter (Fig. 1a, 1b & Video 1). Subsequently, these spherical cells transformed into oval cells (Fig. 1c & Video 2). They then developed surface invaginations (Fig. 1c, 1d & Video 2), which gradually got deeper, transforming an oval-shaped cell into two visibly distinct daughter cells (Fig. 1d & Video 3). These cells then underwent fission, resulting in individual daughter cells. We occasionally saw cells that underwent fission but are still connected by a thin membrane tether (Video 3).

**Fig. 1.**
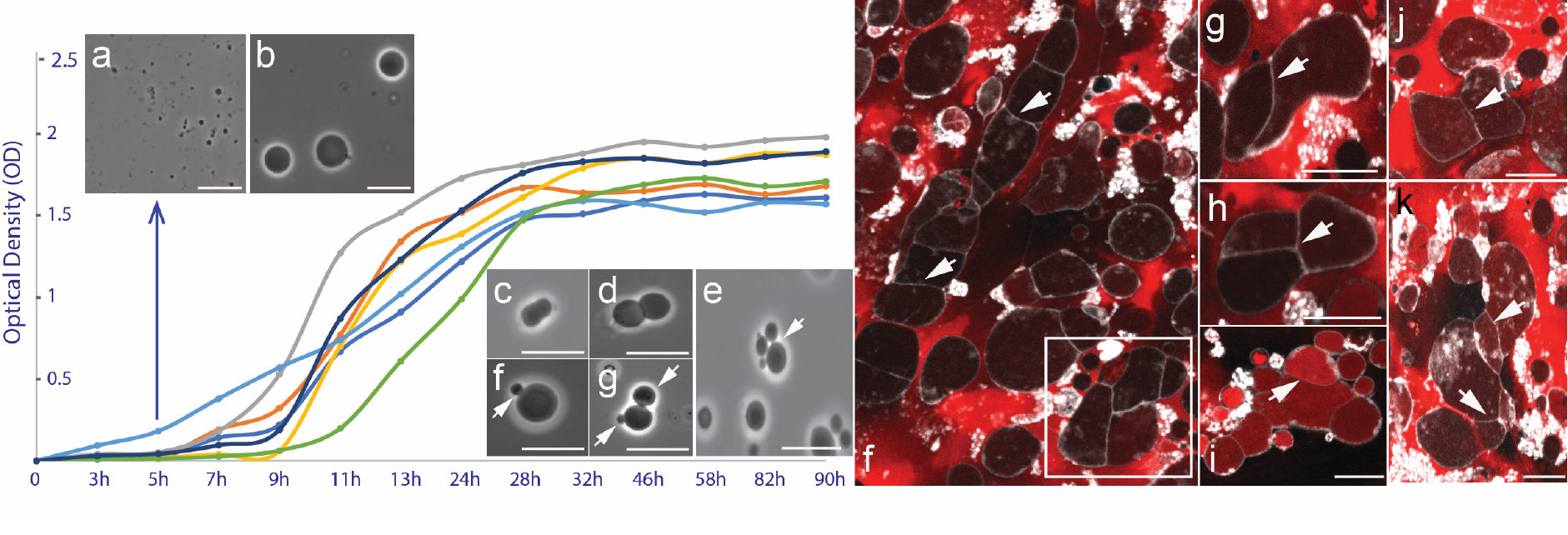
Growth characteristics of *EM-P* in batch culture. *EM-P* exhibited a typical sigmoid growth curve with an initial phase of slow growth rate (lag-phase), a log-phase with exponentially increasing cell numbers and a gradual decline in the growth rate during the stationary-phase. The X and Y-axis in the plot indicate the time in hours (h) and optical density. Images a & b shows typical morphologies of cells in early and late lag-phase. Images c-i show early log-phase cells that seems to be reproducing either by binary fission (c, d and Video 1-3) or budding (f-e, arrows). Images f – k show cells in media isotonic to 7%DSS but devoid of salt (but contain sucrose, BSA and DNA, to maintain appropriate osmolarity). Morphologies of cells under these growth conditions is entirely random. Arrows in the images point to membrane septs formed at random places. Scale bars: 5μm.

One key aspect of the cell cycle is the fission of the parent cell into metabolically viable daughter cells (3). Archiving cell division without the aid of molecular biological processes has been the primary focus of multiple studies and is of crucial importance in enhancing our understanding of most primitive cell replication processes (27) and building autonomously replicating synthetic cells (3, 28). Previous studies archived fission either by mechanical means (29), by encapsulating protein involved in cell division (30), or by an uncontrolled splitting of LVs by gradually increasing the size of the LVs to the point of instability. Given *EM-P* was grown under static conditions as a thin film attached to the base of the culture flask, we do not presume that the physical stimulus required for splitting the parent cell into two daughter cells (binary fission) originated out of the cell.

In accordance with our proposition, most cells in the log-phase were observed to be in a constant state of surface oscillations/deformations (Video 1-3, constantly shivering cell surface). A similar surface instability was theoretically predicted in liquid droplets (31) and experimentally demonstrated in LVs packed with vibrating nanoparticles (32). Such surface deformations in droplets and LVs are shown to have been caused by the Brownian movement of the intracellular molecules and their interactions with the cell membrane. This constant movement of intracellular constituents is also thought to have provided the energy required to split the cell into two daughter cells (32). In tune with these studies, we observed DNA in a constant state of movement (video 4) in *EM-P*. In a normal bacterial cell, DNA forms highly organized and condensed into a structure like a nucleoid or attached to the cell membrane via a cation-mediated salt-bridge (33, 34). The formation of DNA-DNA and DNA-membrane complexes is known to have been mediated by divalent cations, which neutralize negative charges of the phosphate groups (35). In the case of *EM-P*, no such organizational structure was observed. DNA is distributed evenly throughout the cytoplasm as loose strands (video 4). This suggests little interaction between individual DNA strands or DNA and cell membranes. The absence of such structures and high concentration of intracellular DNA due to uncontrolled genome replication (26), could have resulted in repulsion between DNA and membrane. This constant movement of DNA within the cell could have led to surface instability and ultimately to the fission of a parent cell. In addition to the movement of intracellular constituents, similar surface deformations were also known to have been caused by the action of proteins involved in cell division (30).

To understand if the surface deformation were a result of the Brownian movement of intracellular constituents or due to the specific action of some intracellular protein, we treated the cells with bactericidal chemicals like glutaraldehyde and formaldehyde (36). These compounds were known to denature and polymerize proteins with little influence on the biophysical properties of DNA (no influence of on DNA under the experimental conditions of our study)(37). Treatment of cells either with glutaraldehyde and formaldehyde or with 3- methoxybenzamide (38) did not resulted in a static cell. 3-methoxybenzamide inhibits the activity of FtsZ, the tubulin homologue in prokaryotes known to play an important role in regulating bacterial cell morphology and reproduction (39). Moreover, culturing of cells in media containing 3-methoxybenzamide, did not affect the growth characteristics or the morphology of the cells (Fig. S3). These results suggests that innate physiochemical forces rather than the canonical molecular biological processes mediated the cell fission. Similar electrostatic repulsion between old and newly synthesized copies of the genome is recently shown to play an important role in genome segregation and fission of bacterial cells (40).

Together with the Brownian motion of intracellular constituents, surface deformations could also have been caused by the transient binding of cations onto the membrane surface (41) or due to the differences in the osmolarities and composition of molecules between the cytoplasm and the growth media (42–44). In the case of *EM-P*, the osmolarity of the cytoplasm was a result of high packing densities of intracellular constituents (organic in nature)(42). While the osmolarity of the surrounding media was primarily a result of inorganic salts (7%DSS) added to the growth media. To understand the influence of this process on surface instability and cell fission, we transferred cells from 7%DSS-TSB into an isotonic growth media. Unlike 7%DSS-TSB, this media’s high osmolarity resulted from high concentrations of organic compounds like sucrose, glycerol, DNA, and proteins. The presence of these organic compounds could have resulted in similar conditions on either side of the cell membrane. Apart from not observing surface deformations, cells in organic carbon-rich media grew into random morphologies (Fig. 1 f-k). Segregation of the parent cell into two daughter cells did not happen by binary fission. We instead observed random growth of membrane (Fig. 1 f-k), ultimately compartmentalizing single parent cells into multiple daughter cells. Transfer of log-phase cells between these two different media compositions resulted in a reversible change in the cell morphology. Cell surface deformations in 7%DSS- TSB can also be halted by osmotically equilibrating cytoplasm with the surrounding media by treating the cells with membrane pore-forming compounds like Triton X100. These results suggest that cell division in *EM-P* was influenced by the environmental conditions (media composition) and the biophysical properties of cell constituents rather than the action of specific proteins within the cell (45).

Depending on the size of the daughter cells, cells in this growth stage appear to be reproducing either by budding (Fig. 1e-g) or by binary fission (Fig. 1c&d, & 2). Despite numerous repetitions, no uniformity was observed either in the sizes or number of daughter cells formed from a single *EM-P* cell (Fig. 1, 2, S4 & S5d). This asymmetric cell division and variability in the sizes of the daughter cells was theoretically predicted to happen in LVs made from compositionally diverse lipid species (46). Most previous attempts at understanding the biophysical properties of protocells were conducted on LVs of simple chemical composition. Usually made from one to three lipid species(5, 42). In such LVs, the physical properties of vesicle membranes, like elasticity and spontaneous curvature, remain homogeneous along their surface (46). In contrast, live cell membranes comprise numerous lipids species, pigments, quinones, and membrane proteins (47). This diversity results in the formation of lipid-lipid (48) and lipid-protein (49) nanodomains. The physical properties of these nanodomains are dependent on the nature of the interactions between these components (50). Hence unlike the LVs, live cell membranes are considered to be an assembly of nanodomains that differ in their physical properties, i.e., their physical properties like elasticity and curvature remain non-homogenous throughout their surface (46) (Fig. 2a & 2b). This lateral heterogenicity in membrane physical properties, together with the amount of excess membrane, is theoretically predicted to result in cell division resembling budding (46).

**Fig. 2.**
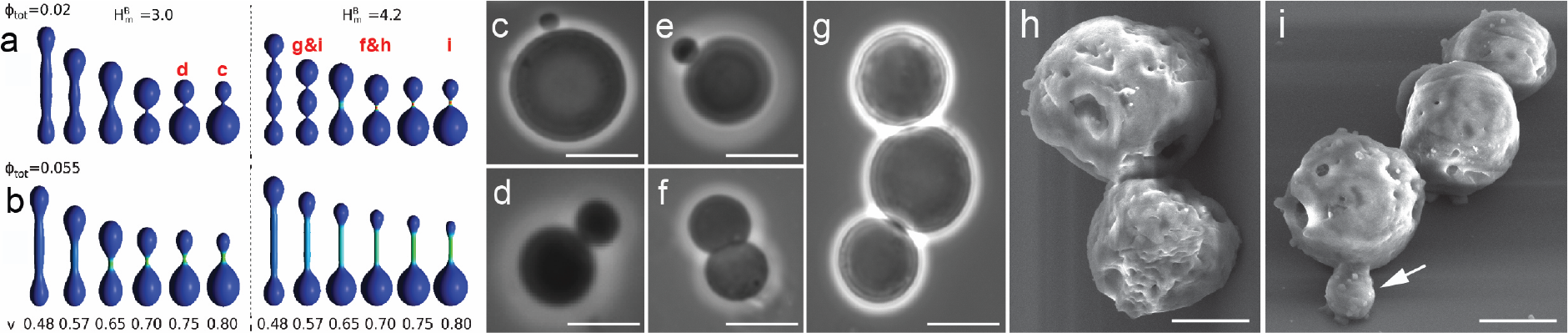
Asymmetric division of *EM-P* cells. Images a & b show theoretically predicted morphologies of vesicles with lateral membrane heterogeneity (originally published by Bobrovska *et al*., 2013)(46). v is the ratio between the actual volume of a vesicle to the volume of a vesicle in its spherical form, H^B^_m_ is the bending rigidity of the anisotropic component (lipid-lipid or lipid-protein nanodomains within the membrane), and Ф_tot_ is the concentration of the membrane component that differed in its intrinsic membrane curvature. Images c-i show, phase-contrast (c-e) and SEM (h & i) images of *EM-P* cells undergoing a similar method of reproduction. Similarities between *EM-P* cells and predicted morphologies is indicated in image a (letters in red). Scale bars: 10μm (c-e) & 2μm (h & i).

Asymmetric cell division and the lack of uniformity in the sizes of the *EM-P* daughter cells can be explained by these theoretical predictions (46). The excess lipid synthesis and diversity of lipid species in *EM-P*’s cell membrane (47) could have created a state of low V (below 1) and H^B^_m_-values (a measure of heterogeneity in membrane physical properties) (Fig. 2). The V-value of the cell is defined as the ratio of the actual volume of the cell (V) and its volume in its spherical form (V_s_), expresses as reduced volume (V; V=V/V_s_). A spherical cell with no excess membrane has V-values of 1. Among all the cell morphologies, spherical cells have the lowest volume compared to the surface area. The presence of excess membrane, as observed in *EM-P* (Fig. S1), increases the cell’s surface area. This, in turn, results in an increased cell volume. This disproportional increase in the cell volume in relation to the cytoplasmic volume results in the transformation of spherical *EM-P* cells into a plethora of non-spherical morphologies. A large difference in the volume of the cell (∼ membrane surface area) and the volume of the cytoplasm results in low V-values (Fig. 2). On the other hand, a smaller difference between these two parameters results in high V-values (closer to 1). Despite the excess lipid synthesis in *EM-P*, most lipids within the log-phase cell were not incorporated into the cell membrane and remained within the cytoplasm as lipid globules (Fig. S1a and S5). This resulted only in a slight elongation of the cells (V-values below but close to 1) and facilitated reproduction by budding (Fig. 2). The diversity in the sizes of the daughter cells can be explained by the regional differences that are to be expected for a bacterial cell, i.e., regional differences in the H^B^_m_-values. To our knowledge, ours is the first study to link the theoretically predicted behavior of LVs with division in live cells (Fig. 2).

During the late log-phase, most cells developed filamentous extensions (video 5) or intracellular vesicles (Fig. 3-6). These structures could have formed due to the gradual transfer of lipids from the lipid globules to the cell membrane (Fig. S5-S7). This transfer of lipids is evident from the absence of lipid globules in stationary-phase cells. This influx of lipids, together with a reduced growth rate (∼slow rate of cytoplasmic volume increase) (Fig. 1) of stationary phase cells, could have resulted in a state of the excess cell membrane (Fig. 3 & S1c-f). The formation of hollow filamentous or intracellular vesicles can be explained as a mechanism for accommodating this excess membrane. In accordance with this inference, cells kept in a continuous log-phase by transferring them to fresh media every few hours did not develop either intracellular or extracellular membrane extensions (Fig. S6 & S7). Like the log-phase cells, these cells continued to reproduce by binary fission (Fig. 1, S5 & S6g).

**Fig 3.**
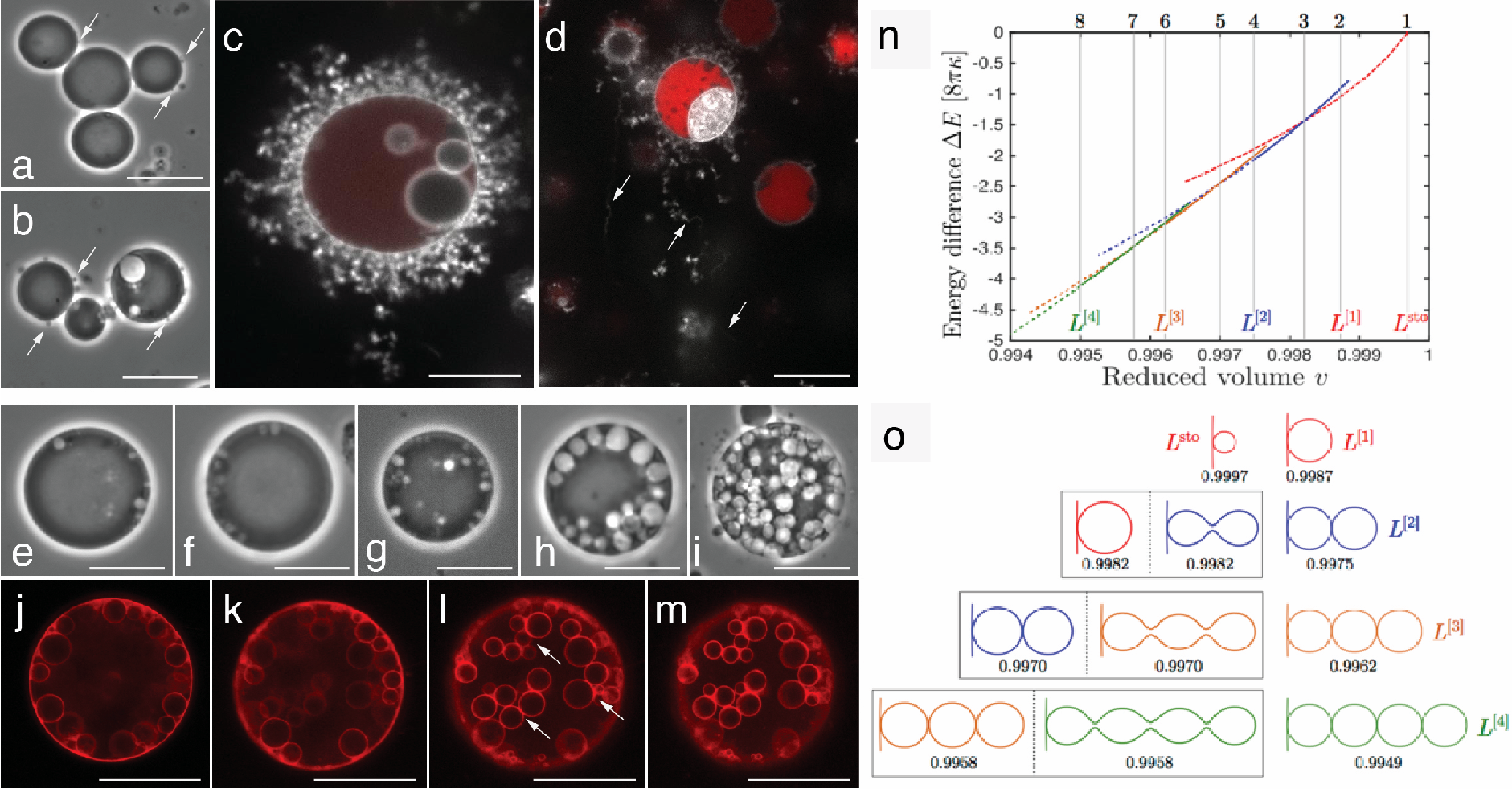
Comparison between theoretically predicted steps in filament formation to observation in *EM-P*. Images a-m show images of *EM-P* cells from different growth stages with external (c-f) or internal membrane (e-m) structures that were formed in a pattern similar to what was theoretically predicted (n & o). Arrows in images a & b point to late-log phase *EM-P* cells with tiny membrane extensions. Image-c shows the cell surface covered with short filamentous extensions. Image - d shows cells transferred to 15%DSS with longer filamentous extensions. Cells in these images were stained with universal membrane stain, FM^TM^5-95 (white), and DNA stain, PicoGreen^TM^ (red). Images e-i show sequential stages observed in the development of intracellular vesicles. Images e-I show vesicles forming at the periphery of the cells before their detachment and migration into the cell. Images j-m show optical sections of a Nile red-stained *EM-P* cell at different planes. Arrows in these images point to the interconnected nature of these vesicles, similar to L(^3^) and L^(4)^ in image-o. Images n & o were originally published by Liu *et al.,* 2015(51). Image-n shows the energy landscape of different equilibrium shapes as a function of reduced volume (v). ΔE indicates deflation-induced (or excess membrane-induced) reduction in the membrane bending energy. Image-o shows the shapes of filamentous extensions corresponding to eight vertical lines in image-n. (please refer to Figure 5 in Liu *et al.,* 2015(51) for further information). Scale bars: 10μm.

Moreover, we also observed the presence of lipid globules within these cells (Fig. S7). The reason for this slow transfer of lipids from the cytoplasm to the cell membrane in actively growing cells is currently unclear.

The formation of filamentous extensions (Fig. 3a-d) and intracellular vesicles (Fig. 3e-i) did not happen in a random manner. These structures were observed to have formed in a sequence of steps previously reported from LVs (Fig. 3 n&o)(51). When LVs are placed in a hypertonic solution, they undergo a step-by-step morphological transformation to reach a thermodynamically stable state (52). The morphologies of such LVs are determined by the osmolarity of the media and by the rate of osmotic shrinkage. The gradual step-by-step increase in the osmolarity of the media resulted in a slow transfer of lipids from the LV membrane to the filamentous structures. From a thermodynamic perspective, this slow transfer of lipids will lead to the formation of hollow bud-like structures from multiple sites on the cell surface (51). Whereas a sudden increase in osmolarity resulted in a rapid shrinkage of the LVs (i.e., a relatively faster rate of lipid transfer), which was shown to thermodynamically favor the formation of fewer but longer filamentous extensions (Fig. 3n&o).

Like in LVs, the sequential steps involved in the formation of membrane extensions can be explained by the differences in the growth rates of log and stationary phase cells and by the rate of lipid transfer from the LV into the membrane extensions. We observed lipids accumulated within the cytoplasm during the log-phase as lipid droplets (Fig. S1). We noticed a slow rate of lipid transfer between these droplets and the cell membrane during the log-phase cells (Fig. S7). This slow transfer of lipids to the cell membrane and the high growth rate (∼ rate of cytoplasmic volume increase) and nearly spherical morphologies suggest that cells maintained a close match between the cell volume and surface area of the cell membrane, i.e., less membrane was available for forming hollow filamentous extensions. This led to the slow transfer of lipids from the cytoplasm to the membrane resulting in the formation of tiny hollow membrane buds from multiple sites on the cell surface (Fig. 3a, b, and e-h). Complete transfer of lipids from lipid droplets to the cell membrane during the late-log and early stationary-phase, resulted in a sudden influx of lipids from the cytoplasm into the cell membrane. Together with the reduced growth rate during the stationary phase, could have energetically favored an increase in the length of the existing membrane extensions (both internal and external Fig. 3d & 3l-m) rather than forming new membrane invaginations on the cell surface. Transfer of late log-phase cells from media with 7%DSS to 15%DSS to create a sudden state of lipid excess led to osmotic shrinkage of cells and abrupt increase in the membrane area. In accordance with previous observations on LVs(51), these cells were observed to have formed longer and fewer filamentous extensions (video 6) without the initial appearance of multiple membrane buds on the cell membrane. These results suggest that growth rate and environmental conditions played an important role in determining the formation and morphology of filamentous extensions (Fig. 3).

All the above discussed morphological changes like the budding and formation of external and internal membrane structures is associated with phase-separation of membrane into more fluid, liquid-disordered (L_d_) and more rigid, liquid-ordered (L_o_) phases. This appearance of distinct phases in the cell membrane and associated formation of external and internal membrane extensions can be explained by the model proposed by Julicher & Lipowsky (53, 54). It was hypothesized that phase-separated membranes minimize interfacial energy at the phase boundary to attain a thermodynamically stable state. They do so by forming an out- of-plane invagination (bud formation), gradually transforming the softer L_d_-membrane into a bud. In accordance with this theory (53, 54), the formation of buds in *EM-P* is associated with the appearance of a distinct membrane phase – more fluid, L_d_ and more rigid L_o_ membrane (Fig. 4). L_d_ membrane initially appeared as small patches on the cell surface, less than 180nm in diameter (Fig. 4a-c). As the area of the L_d_ membrane increased, it underwent budding due to an increase in the line tension at the phase boundary (55)(Fig. 4a, 4d, 4g & S8). However, we observed some deviations from this expected behavior. First was the difference in the direction of budding. Contrary to the predictions (53), we observed L_d_-membrane underwent a negative (into the cell) rather than positive (out of the cell) curvature (Fig. 4g). Second, despite its higher bending rigidity and unfavorable thermodynamics, we also observed a more rigid L_o_-membrane undergoing budding. All the membrane structures extending out of the cell are exclusively composed of L_o,_ and all the membrane structures extending into the cell were composed of L_d_ membrane (Fig. 4d-f, & arrows 4g, cell within the highlighted region).

**Fig. 4.**
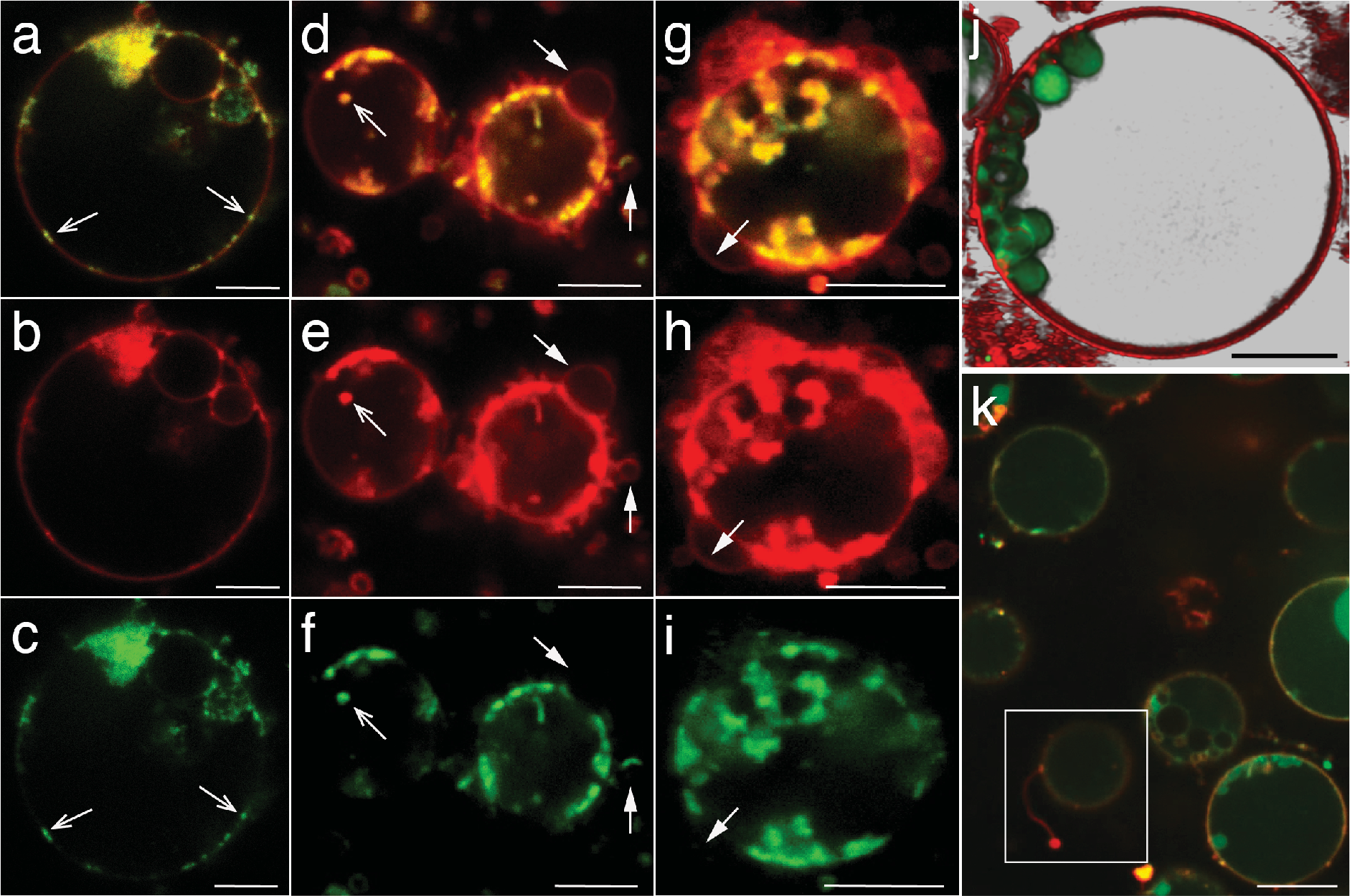
**Membrane phase-separation in *EM-P*. Membrane phase-separation in *EM-P*** Cells in these images were stained with FM^TM^5-95 (all membrane, red) and *FAST*^TM^ DiI (L_d_- membrane, green) and imaged either using a super-resolution STED microscope (a-j) or a spinning disk confocal (k). Images a-c, d-f, and g-i are images of the same cells in different confocal channels. Open arrow in a-f, point to small patches of L_d_-membrane on the cell surface (≈ 180nm, beyond the diffraction limit of the conventional microscopes) (a-c). Closed arrows in d-i: L_o_-membrane extensions out of the cell. Image j is the 3D-rendered image of intracellular vesicles composed of L_d_-membrane within the cell. A series of optical sections of the stacks is shown in Fig. S8. Image k shows *EM-P* cells with filamentous extensions, as shown in Figure 3c, entirely composed of L_o_ membrane. Scale bars: 10μm (a-i) & 5μm (j & k).

The direction and physical nature of a membrane that underwent budding in *EM-P* could be explained by its lipid composition (56) and high molecular crowding of the cytoplasm (57). Bacia *et al.* studied the influence of membrane composition on budding (56) and reported that the membrane sterols composition influences the direction and nature of the membrane that undergoes budding. Despite higher bending rigidities, this study showed that some sterols could preferentially induce the budding of a more rigid L_o_ membrane instead of L_d_ membrane. Prokaryotes lack sterols like cholesterol and their derivatives but possess functional cholesterol analogs, like hopanoids (58). The properties of these compounds in inducing membrane phase separation and budding were rarely investigated. Apart from sterols, invagination of a more rigid L_o_ membrane was also reported to happen when LVs (52) experience a state of an excess membrane (during the osmotic shrinkage in LVs or slow growth rate in protoplasts), possibly due to an increase in the spontaneous curvature of the L_o_ under such conditions (59).

The nature and direction of the membrane invagination could also have been determined by the differences in the molecular crowding between the cytoplasm and the growth media (57). Studies showed that when the aqueous core (cytoplasm) undergoes phase-separation into regions of higher and lower densities, results in a spontaneous membrane phase-separation (57). In such two-phase LVs, the L_d_ and L_o_ phases of the membrane were observed to have been associated with cytoplasmic regions of higher and lower densities, respectively. The mechanism behind this preferential association is currently not well understood. Nevertheless, this could explain the selective invagination of the L_d_ into the cell given the higher densities of cytoplasm (∼17-35 wt% molecules) and L_o_ membranes into and out of the cell due to the comparatively less dense growth media (∼6-8 wt% molecules) (Fig. 4). TEM images of cells shown in Fig. 6, show these differences in the packing densities between the cytoplasm and surroundings (60).

The live cell membranes are thought to undergo phase-separation, a process proposed to profoundly influence cellular functions (48). Despite these propositions, the existence of distinct phases within the cell membrane has been a point of contention among researchers (61). To the best of our knowledge, ours is among the very few studies that provide visual evidence of membrane phase-separation in live cells (Fig. 4)(62).

The formation of filamentous extensions and intracellular vesicles (Fig. 3) led to two different reproduction methods. In short, *EM-P* reproduction either by forming external (Fig. 5) or intracellular (Fig. 6) daughter cells. Sequential steps involved in forming external and internal daughter cells are shown in videos 7-11 and 12-18, respectively.

**Fig. 5.**
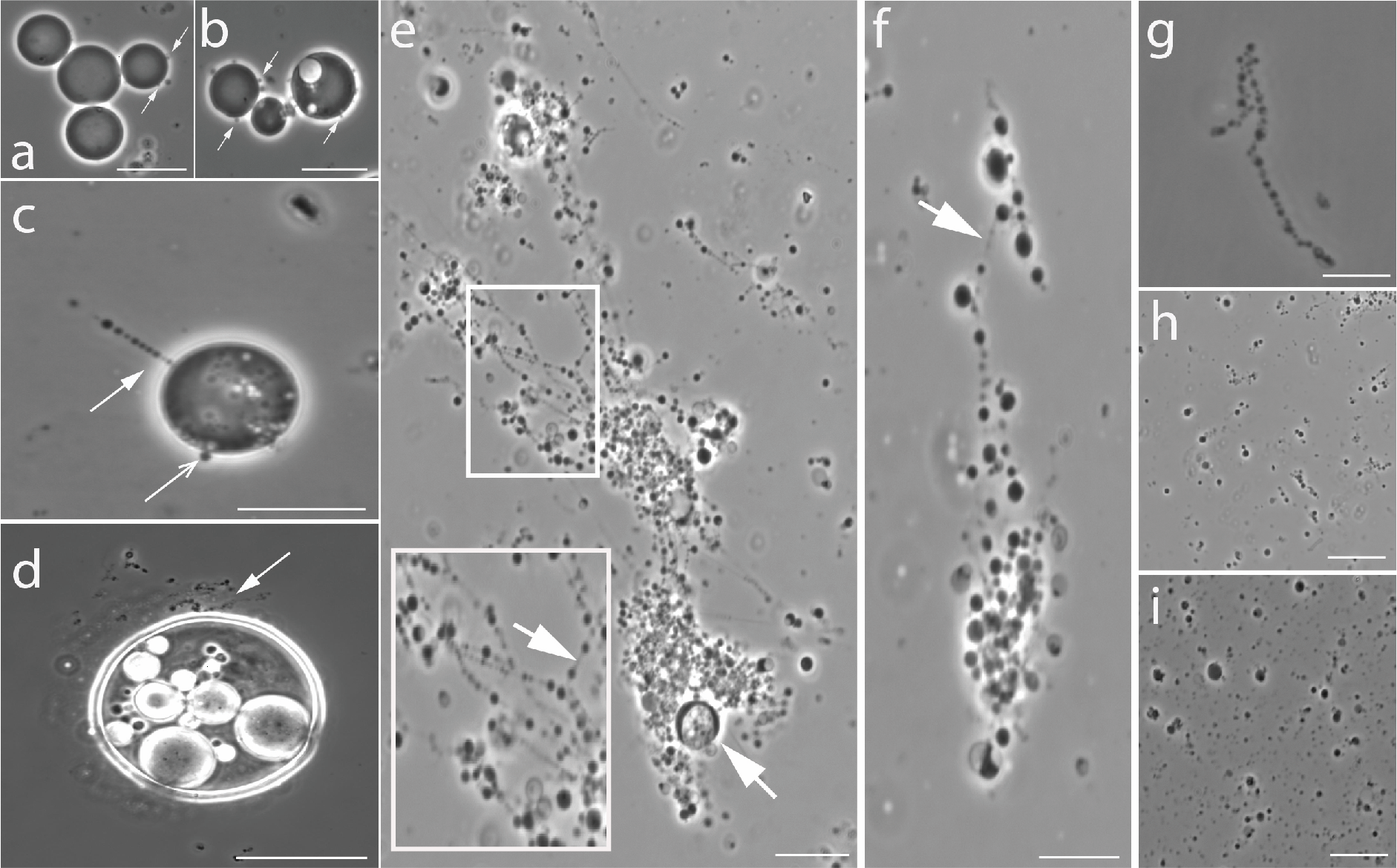
Life cycle of *EM-P* reproducing via formation of extracellular daughter cells. Image a & b show a late log phase cell with bud like structures on *EM-P* cell surface. Image- c show a late log phase cell with open arrow pointing to surface buds and a closed arrow pointing towards filamentous extensions. Image-d shows a cell from a subsequent growth stage cells with multiple filamentous extensions (also see video 5). Images e-f show formation of daughter cells as “string-of-beads” attached to a parent *EM-P* cell (boxed region) (also see video 6-8). Images f-i show the detachment of these strings of daughter cells and their subsequent fragmentation into individual daughter cells (also see video 9-11). Scale bars: 10μm (a-i).

During the stationary growth phase, we observed cytoplasm transfer from the parent cell into filamentous extensions (Fig. 5c & Video 7-9). Consequently, the hollow tubular filaments transformed into a “string-of-individual daughter cells” (Fig. 5c-g & Videos 8&9). The transformation of hollow filamentous extensions into string-of-individual daughter cells could be explained by two different mechanisms. One could have been the spontaneous transformation of tubular vesicles into a “string-of-beads” morphology to reduce the interfacial area to reach a thermodynamically stable lower energy state(63). A similar transformation of tubular vesicles during osmotic shrinkage when placed in a hyperosmotic solution (identical to *EM-P,* video 6) was previously reported by multiple studies (63, 64).

Alternately, the formation of daughter cells could also be explained simply by the lower volume of cytoplasm that was transferred into the filaments compared to the total volume of the filament (51). The excess membrane within these tubular structures should have collapsed and transformed into tethers (Fig. 5 e&f, arrows) connecting adjacent daughter cells.

Lowering the salt content (osmolarity) of the media from 7% to 4 %DSS during the late log-phase prevented the formation of filamentous extensions and transfer of cytoplasm to leading to the formation of daughter cells (Fig. S9). Possibly due to the influx of water these cells we observed these cells to be larger than the cells in the control experiment (Fig. S9). This absence of daughter cells could be explained by the osmotic expansion of cells which could have prevented the cytoplasm transfer from the cell into the filamentous extensions. In comparison, the transfer of cells into media of higher osmolarity (15%DSS) resulted in the osmotic deflation of cells and the formation of longer filamentous extensions with individual daughter cells (Fig. S10). These strings of daughter cells are then detached from the parent cell and fragmented into individual daughter cells in a sequence of steps, as shown in Figure 3F-I (video 9-11).

We also observed the formation of daughter cells within the intracellular vesicles. These daughter cells were formed by a process similar to budding (Fig. 6d, arrow). These buds either detached from the vesicle surface and released into the vesicle as individual daughter cells (Fig. 6f&h & video 14) or grew into a string of daughter cells (Fig. 6e & video 14 – cell on the left). These “string-of-daughter cells” subsequently detached from the vesicle and fragmented (Fig. 6h) into individual daughter cells (video 17). Due to the gradual cytoplasm transfer from parent to daughter cells (Fig. 6d, & 6e), the cytoplasmic content of the *EM-P* cells gradually depleted (Fig. S11). By the late stationary-phase, many *EM-P* cells were hollow with intracellular vesicles (Fig. 6g, S4 & video 14) containing highly motile daughter cells. Daughter cells were released into the surrounding media by rupture of the cell or vesicle membrane (Fig. 6h & Video 12-18). The Reproduction by formation of intra vesicular daughter cells is observed in previous studies (65). Unlike in our study which shows that daughter cells were released into the intracellular vesicles, these studies presumed that hollow vesicles to be daughter cells. To the best of our knowledge, ours is the first study to show the sequential steps involved in the formation of internal daughter cells. The final growth stage of *EM-P* was characterized by the presence of cells with all the above-described morphologies, along with a considerable amount of membrane debris (Fig. S12).

**Fig. 6.**
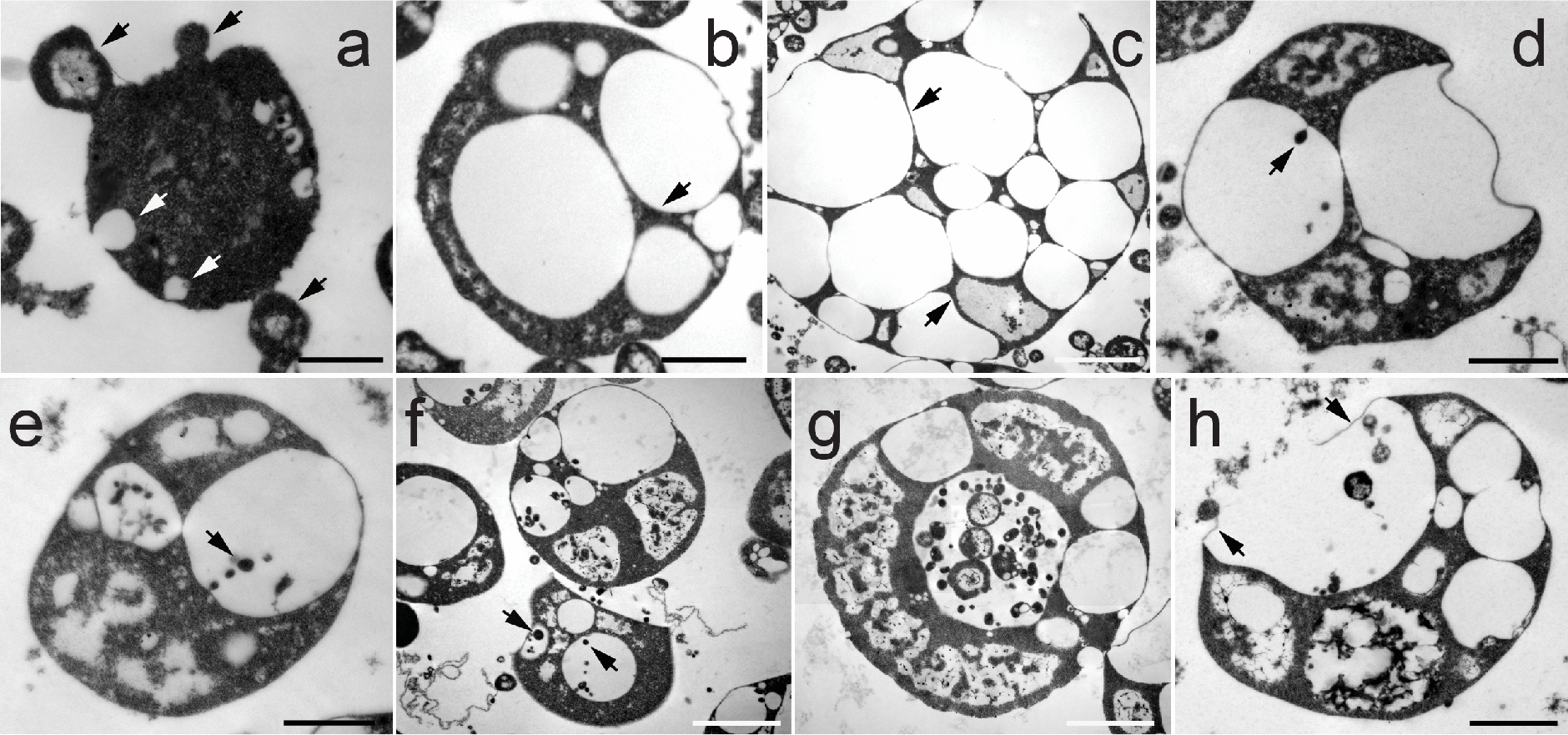
The life cycle of *EM-P* reproducing via the formation of intracellular daughter cells. Images a-h is TEM images of the life cycle of *EM-P* reproducing by the formation of intracellular daughter cells. Image-a show the process by which intracellular (white arrows) vesicles and bud-like structures are formed (black arrows). Closed red arrows in images b & c point to the narrow cytoplasmic region of the cell. Arrows in images d-f points to the formation of individual and string of daughter cells or string of multiple daughter cells. Image-g shows internal daughter cells with intracellular vesicles. Image-h shows the daughter cell release process by rupturing the vesicle membrane (open red arrows). Scale bars: 1μm.

Much of the focus in the above sections was dedicated to understanding *EM-P*’s life cycle from a biophysical perspective. Apart from the overall salinity, the biophysical properties of the cell constituents were known to have been influenced by the nature of the salts in the media. Different cations interact differently with the cell membrane and alter their physical properties (42,66,67). To understand how individual components of DSS influenced the morphology and life cycle of *EM-P*, we replaced DSS in the growth media with an equivalent concentration of pure salts like KCl and MgCl_2_ (6%w/v). When grown in TSB-7%KCl, cells reproduced predominantly via the formation of intracellular daughter cells (Fig. S13). Cells in these incubations are larger than cells grown in DSS. In contrast, cells grown in TSB- 6%MgCl_2_ were significantly smaller than those grown in equivalent concentrations of DSS or KCl (Fig. S14, video 19 & 20). The diameter of these cells was restricted to 0.5-2μm and reproduced by forming extracellular daughter cells (Fig. S14). The nature of interactions between cations and phospholipid head groups(42) can explain these morphological differences. Earlier studies reported that monovalent cations like potassium increase the membrane’s fluidity, whereas divalent cations like magnesium stiffen the membrane by reducing its fluidity(66). This increase in membrane fluidity could have thermodynamically facilitated the easier expansion of cell membranes in incubations containing DSS and KCl (42), resulting in relatively larger cells. Increased stiffness of the membrane in the presence of MgCl_2_ could have prevented the easier expansion of the cell membrane resulting in the formation of smaller cells (Fig. S14). The reasons for the reproduction predominantly via extracellular daughter in the presence of MgCl_2_ and via intracellular daughter cells in the presence of KCl are currently unclear. Nevertheless, a similar phenomenon of intracellular vesicle formation in the presence of KCl was observed previously in LVs (42). *EM-P* reversibly changes its morphology when transferred between media of different salt compositions.

Apart from the salinity, one other characteristic feature of the coastal marine environments is tidal activity. Given the moon’s proximity during the Archaean Eon, it is thought that coastal marine environments of early Earth could have experienced high-intensity wave action (68). To understand the influence of such physical forces on *EM-P’s* reproduction, we grew cells on an orbital shaker. Under these growth conditions, *EM-P* reproduced exclusively by a process resembling binary fission, like the cells in the early log-phase (Fig. 1c-e). None of the above-described morphologies, like large cells with intracellular vesicles or strings of daughter cells (Fig. 5&6), were observed in these incubations. However, we observed the re- appearance of filamentous extensions and intracellular vesicles when we transferred late log- phase cells from an orbital shaker to a static incubator. These observations suggest that reproduction by binary fission is a result of culturing the cells under non-static conditions.

Transfer of *EM-P* between media containing different salts or from an orbital shaker to static conditions led to a reversible change in cell morphology.

A considerable amount of membrane debris (Fig. S12) and extracellular DNA (eDNA) was observed during all growth stages (Fig. 7 & S15). These observations imply that *EM-P’s* reproduction is leaky, resulting in a significant loss of intracellular constituents. DNA could have been released into the surroundings when the cell underwent lysis to release the daughter cell (Fig. 6h, S5 & video 14-18) or was a result of cells undergoing explosive lysis during all growth stages (Fig. S16 & video 21). To our knowledge, such an explosion of bacterial cells was not reported previously. However, a similar rupture of LVs by two different mechanisms is currently known (69, 70). This lysis of *EM-P* could have been caused either due to osmotic pressure created by Reactive Oxygen Species (ROS)(70) or by interactions between DNA and lipid head groups (69). To understand the role of ROS in *EM- P* lysis, we grew *EM-P* under anaerobic conditions in media devoid of terminal electron acceptors (fermentation conditions). ROS are generally produced by the activity of ETC (71). Given the absence of terminal electron acceptors and the high catalase activity of *EM-P*, we presume low ROS production and their effective neutralization under anaerobic conditions.

**Fig. 7.**
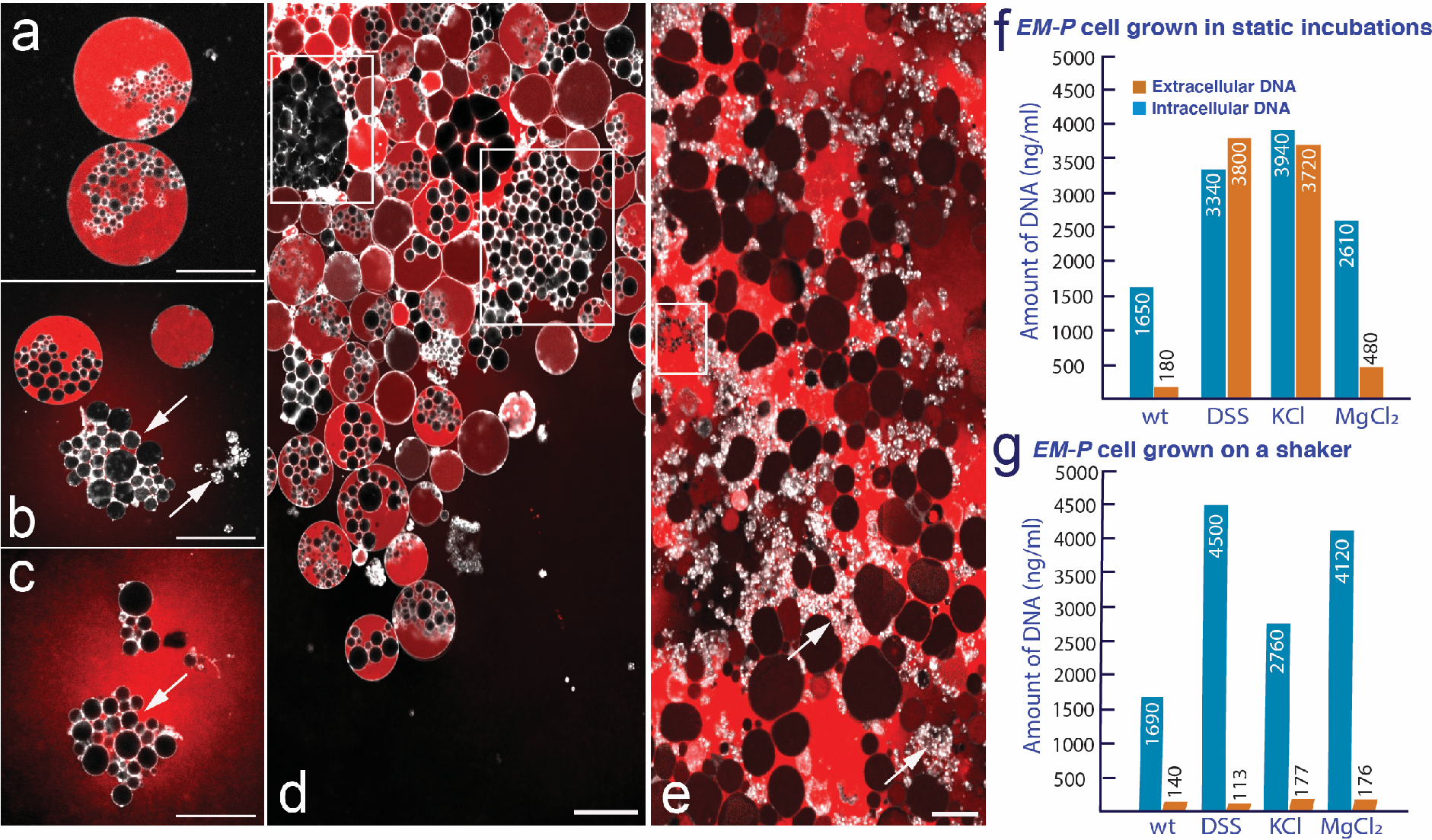
Leakage of cell constituents during *EM-P*’s reproduction. Image a-c shows lysis of *EM-P* cells and release of intracellular DNA (red). Cells in these images were stained with FM^TM^5-95 (membrane, white) and PicoGreen^TM^ (DNA, red) and imaged using a STED microscope. Image-d shows a similar process in the biofilm from the static incubations. Highlighted regions in this image show hollow intracellular vesicles released by the lysis of cells. Image-e show late log-phase *EM-P* cells with considerable amount of extracellular DNA, hollow intracellular vesicles (boxed regions), and membrane debris (arrows). Scale bars: 10μm. Image-f & g show the results of the quantification of DNA leakage during *EM- P’s* late log growth stage (Figure S15, for all growth stages) grown under static conditions and on an orbital shaker respectively. wt – wild type (*Exiguobacterium strain M* with a cell wall); DSS, KCl & MgCl_2_: *EM-P* grown in TSB amended with 7% of the respective salts. Data in images f & g represent mean values of biological repetitions from different batches of *EM-P*’s inoculum’s (n=5).

Given the possibility of photoinduced ROS production (72), all bottles were incubated in the dark and imaged them using a phase-contrast microscope at fewer intervals (48h – 72h).

There were no significant differences in the amount of eDNA between the aerobic control and anaerobic incubations (Fig. S16). Apart from the eDNA, we also observed all the end products of cell lysis, like membrane debris, in these incubations. These results suggest that cell lysis is a natural part of *EM-P*’s cell cycle rather than an artifact of the imaging process.

The efficiency of *EM-P* reproduction also depended on the environmental conditions. We observed less leakage of cell constituents when cells were grown in the presence of MgCl_2_, compared to other salts (Fig. 7 & S15). Significantly less leakage of cell constituents was observed when *EM-P* was grown on an orbital shaker than under static conditions (Fig. 7 & S15). These observations suggest that the reproductive efficiency of *EM-P*-like cells could be higher in natural environments than in our static incubations. Despite the low reproductive efficiency, *EM-P* batch cultures always resulted in viable daughter cells (Fig. 8, S17 & S18).

**Fig. 8.**
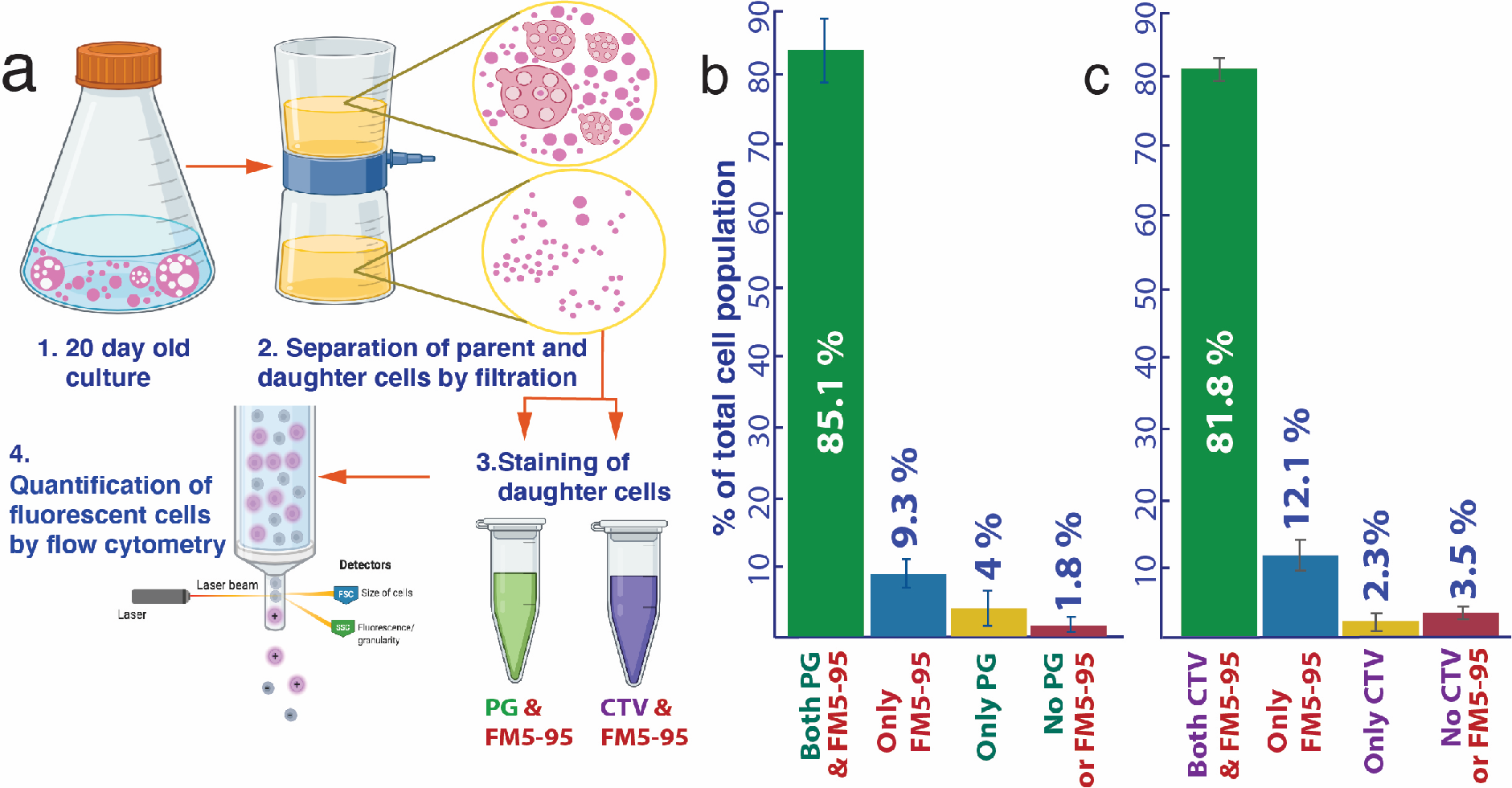
Metabolic viability of daughter cells. Panel-a shows the workflow of the experiment. Image-b shows the results of this quantification of these cells by flow cytometry. The green bar shows the percentage of cells containing both an intact membrane and intracellular DNA (FM^TM^5-95 & PG). The subsequent blue, yellow, and red bars show the percentages of cells that were either deficient in intracellular DNA or an intact membrane or both respectively. Image-c shows the results of this quantification of different cell population flow cytometry. The green bar shows the percentage of cells containing both an intact membrane and intracellular enzyme activity (FM^TM^5-95 & CTV). The subsequent blue, yellow and red bars show the percentages of cells that were either deficient in intracellular enzyme activity or an intact membrane or both respectively. The original flow cytometry plots together with the confocal microscope images of daughter cells were shown in Figures S17 & S18.

Staining and quantifying cells by flow cytometry showed that most daughter cells received DNA from the parent cell and exhibited cytoplasmic activity (Fig. 8). Moreover, inoculating smaller daughter cells after passing through a 0.44μm filter to separate them from membrane debris and other larger cells always resulted in growth, suggesting the viability of daughter cells. The transfer of daughter cells from 7%DSS-TSB to media containing glucose as the sole carbon and energy source always resulted in growth. This suggests that daughter cells received a full complement of genes from the parent cells, which is necessary for synthesizing all the cell components.

Throughout our study, we observed the morphology and method of reproduction was determined by the environmental conditions. Transferring cells from salt rich to sugar rich media (Fig. 1), changing the osmolarities (Fig. S9&S10) or salt composition within the media (Fig. S13&S14), or the incubation conditions (static or stirring) had a significant influence on the cell morphology, reproduction, and reproductive efficiency (Fig. 7&S15). Transfer of *EM-P* between media containing different salts or from an orbital shaker to static conditions led to a reversible change in cell morphology. This suggests that environmental conditions rather than the information encoded in its genome determined all aspects of *EM-P*’s life cycle. Such profound influence of environmental conditions on bacterial cell reproduction has never been reported before. Our study also demonstrates the working with protoplasts rather than lipid vesicles as proxy-protocells and working under natural environmental conditions rather than well-controlled laboratory conditions provides better understanding of the behavior of protocells.

Our current understanding of protocells results from either a bottom-up approach of building a synthetic cell from scratch (2, 3) or a less frequently used top-down approach of transforming extant bacteria into a primitive state by reducing its genome size to a bare minimum (73). The method of reproduction we observed in *EM-P* resembles the reproductive process of minimal cells resulting from both these approaches (73) together with the existing theoretical predictions on protocells (74)(Fig. S19 – S21). Despite these similarities, suggesting protocells reproduced by this process first requires us to prove the existence of such cells on early Earth. To date, protocells remain a theoretical possibility with no definitive proof of their existence. To understand if protocells ever existed on early Earth, we compared the distinctive *EM-P* morphologies with the morphologies of the earliest known microfossils from Paleo-Archaean Eon. In this follow-up study, we demonstrated the remarkable similarities between *EM-P* and microfossils in terms of their morphology, elemental, and isotopic composition (Additional materials). The results of this study provide an independent paleontological conformation of our proposition that protocells inhabiting early Earth reproduced by a process similar to *EM-P* (10).

## Conclusion

Our results suggest that reproduction is an intrinsic biophysical property of cells with an internal metabolism. Even in the absence of canonical molecular biological mechanisms that regulate reproduction, cells can still reproduce efficiently. In such cells, environmental conditions play an essential role in determining their method of reproduction and reproductive efficiency. Despite the long-held notion that protocells reproduced independent of canonical molecular biological processes and environmental conditions influenced protocell reproduction, ours is the first study to demonstrate this process experimentally.

Given the suitability of this process to the environmental conditions of early Earth, its simplicity, and its efficiency, we propose that uni-lamellar or Gram-positive protocells inhabiting early Earth likely reproduced by a similar process.

## Acknowledgements

We want to thank Petra Schwille for her support and scientific input throughout the work and in preparing the manuscript. Gabriella Berthal, and Markus Oster for excellent technical support. We thank the Advanced Light Microscopy Facility at EMBL, Heidelberg, Ulf Schwartz from Leica Microsystems, and colleagues at the departments of Ecological Microbiology (Bayreuth University) and of Cellular and Molecular Biophysics (Max Planck Institute for Biochemistry) for their support throughout the work.

## Competing Interests

The authors declare that there are no conflicts of interest regarding the publication of this article.

## Data Availability Statement

All data, materials, and methods will be shared by Dr. Dheeraj Kanaparthi upon reasonable request.

## Notes

### Competing Interest Statement

The authors have declared no competing interest.

### Summary of Updates

new data

