## Supplementary materials 1 for "The reproduction process of Gram-positive protocells"

**Supplementary results & discussion:**

**Physiological characterization of protoplasts**

*Exiguobacterium strain M* (*EM*) was transformed into protoplasts, as described in the methods section. This transformation of "wild-type" *Exiguobacterium strain M* (*EM*) into its protoplast state (*EM-P*) was evident from the change in morphology from bacillus to spherical morphology. Together with morphology, considerable differences were observed in the physiological behavior of *EM* and *EM-P*. One such difference was the glucose metabolism under aerobic conditions (Fig. S1). We observed the accumulation of fermentation products like lactate, acetate, formate, and ethanol when *EM-P* was grown in well-aerated media containing glucose as a sole carbon source. In contrast, no such formation of volatile fatty acids was observed in *EM* incubations. This suggest that *EM-P* is metabolizing glucose by random metabolic processes irrespective of the environmental conditions.

*EM* and *EM-P* also exhibited differences in catalase activity. *EM* exhibited an expected trend in catalase activity. Catalase activity was observed when the *EM* cells were grown under aerobic conditions (58.7 ± 6.2 U/ml), while little to no catalase activity was exhibited by *EM* under fermentation conditions (3 ± 0.8 U/ml). In contrast, *EM-P* showed catalase activity independent of the growth conditions. A similar level of catalase activity was observed when the cells were grown under aerobic (77.8 ± 5.3 U/ml) and anaerobic (65.3 ± 7.2 U/ml) conditions.

Catalase enzyme is expressed to counter act the reactive oxygen species produced by the electron transport system. Expression of such enzymes is usually repressed during anaerobic growth. This expected trend was observed in wild-type *EM* cells but not in *EM-P*. Together with the differences observed in glucose metabolism, these results suggest a dysfunctional intracellular coordination.

**Supplementary figures:**

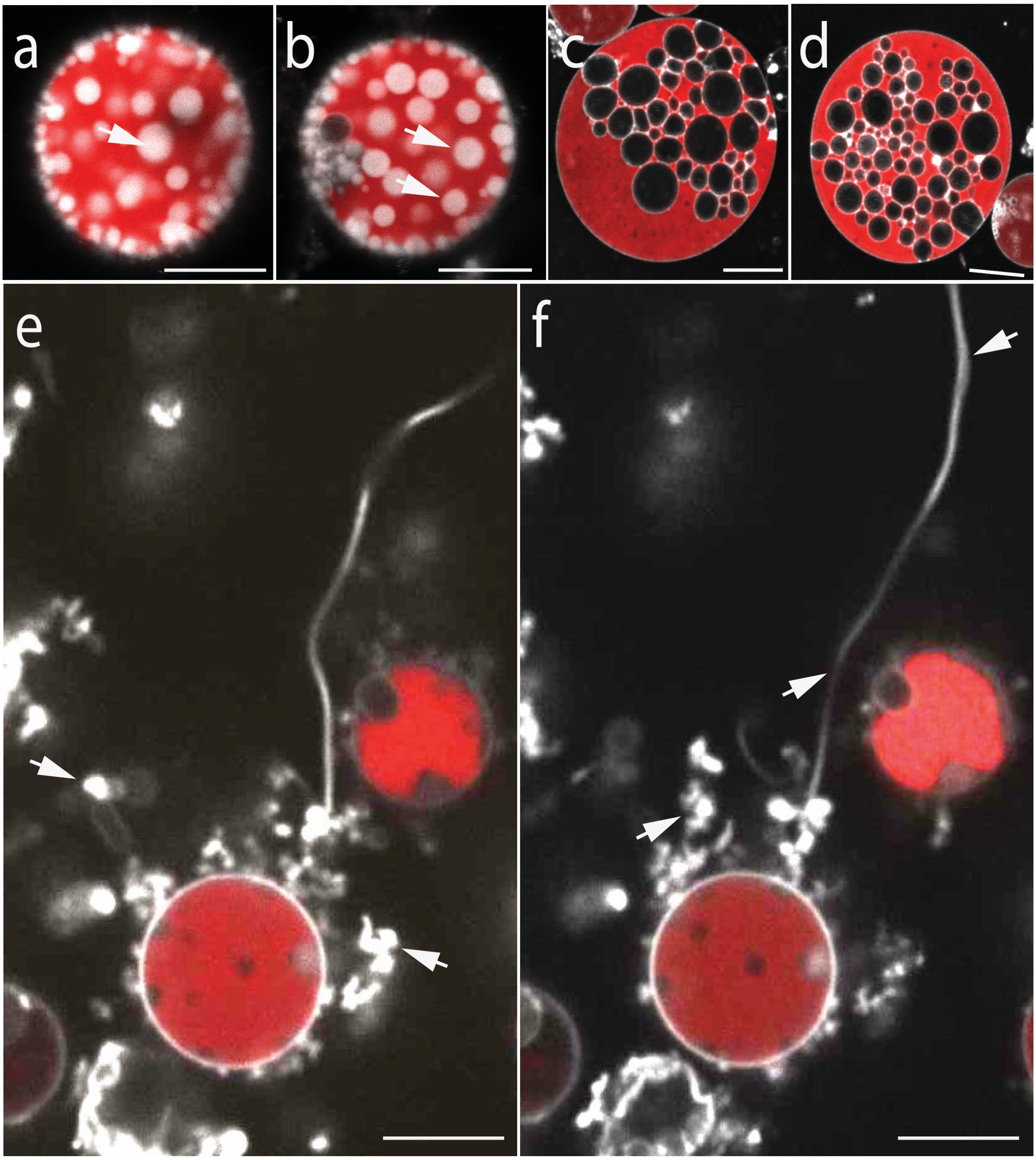


**Fig. S1 *EM-P* cells in different growth stages exhibiting excess cell membrane**

Images a-f are the STED (a-d) and spinning disk confocal (e&f) microscope images of *EM-P* cells. Cells in these images were stained with universal membrane stain, FM™5-95 (white), and DNA stain, PicoGreen™ (red). All cells exhibit excess membrane in the form of lipid globules (a & b, arrows), hollow intracellular vesicles, and filamentous extensions (e & f, arrows). Scale bars: 10µm.

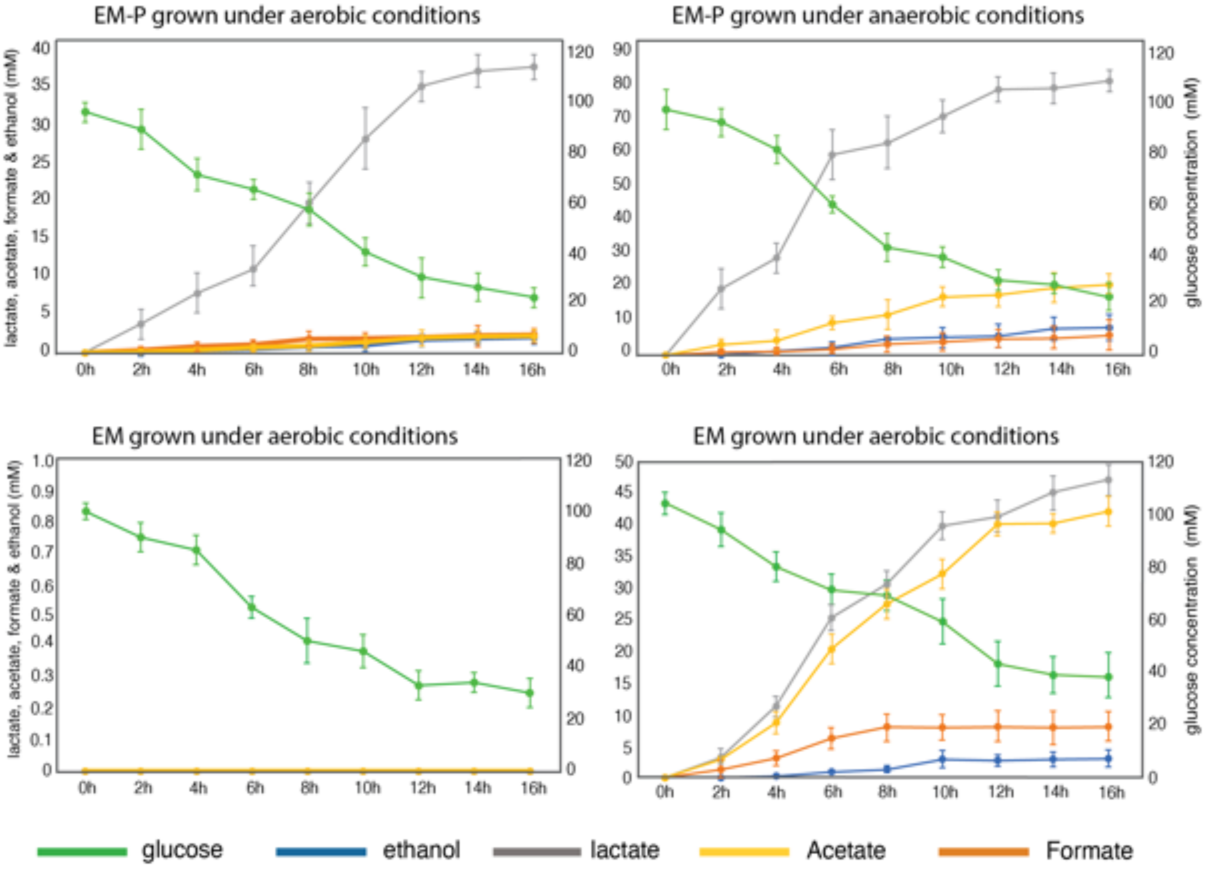


**Fig. S2 Glucose consumption by *EM-P* and *EM* under aerobic and anaerobic conditions**

The plot shows a gradual consumption of glucose and an increase in volatile fatty acids like lactate, butyrate, and acetate during the growth of *EM-P* and *EM*. The X-axis shows time (hours), the primary Y-axis shows the concentration of volatile fatty acids (ethanol, formate, acetate, and lactate), and the secondary y-axis shows glucose concentrations. All the measurements were done in replicates (n=5).

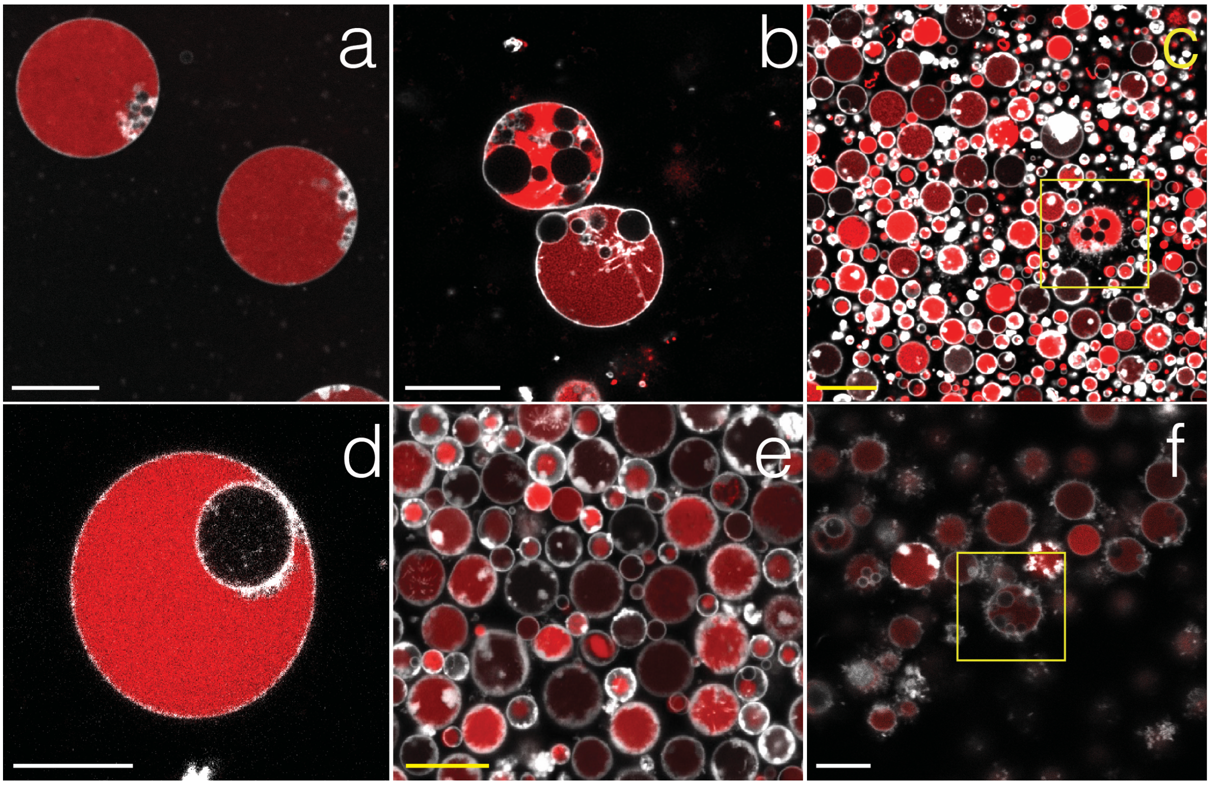


**Fig. S3 Morphological comparison of cells grown in the presence and absence of 3-methoxy benzamide**

Images a-f are STED microscope images of *EM-P* grown in the presence (a-c) and absence (d-f) of 5 mM 3-methoxy benzamide. No significant differences were observed in the morphology or reproduction between these two incubations. Cells in these incubations developed intracellular vesicles and filamentous extensions (highlighted regions in images c & f). Cells in these images were stained with universal membrane stain, FM™5-95 (white), and DNA stain, PicoGreen™ (red). Scale bars: 10µm.

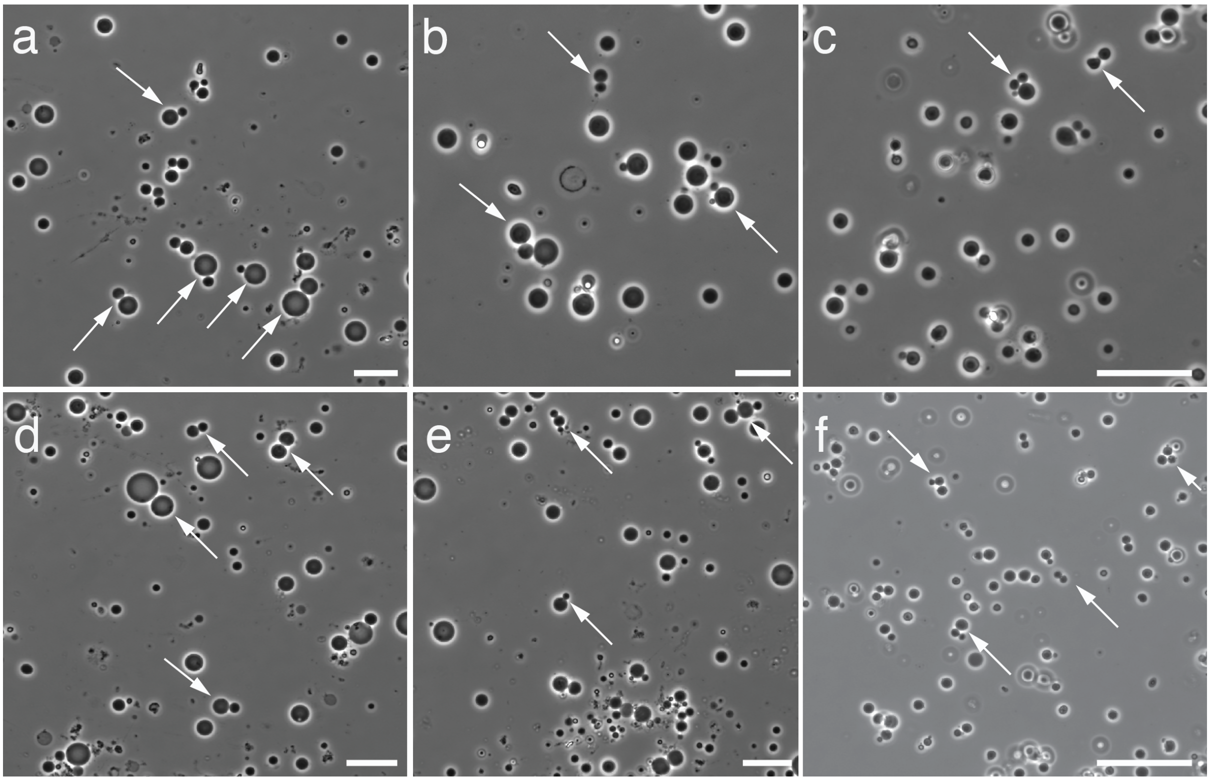


**Fig. S4 Log-phase *EM-P* cells undergoing reproduction**

Images a-f show phase contrast images of log-phase *EM-P* cells that appear reproducing by binary fission or budding. Scale bars: 10µm.

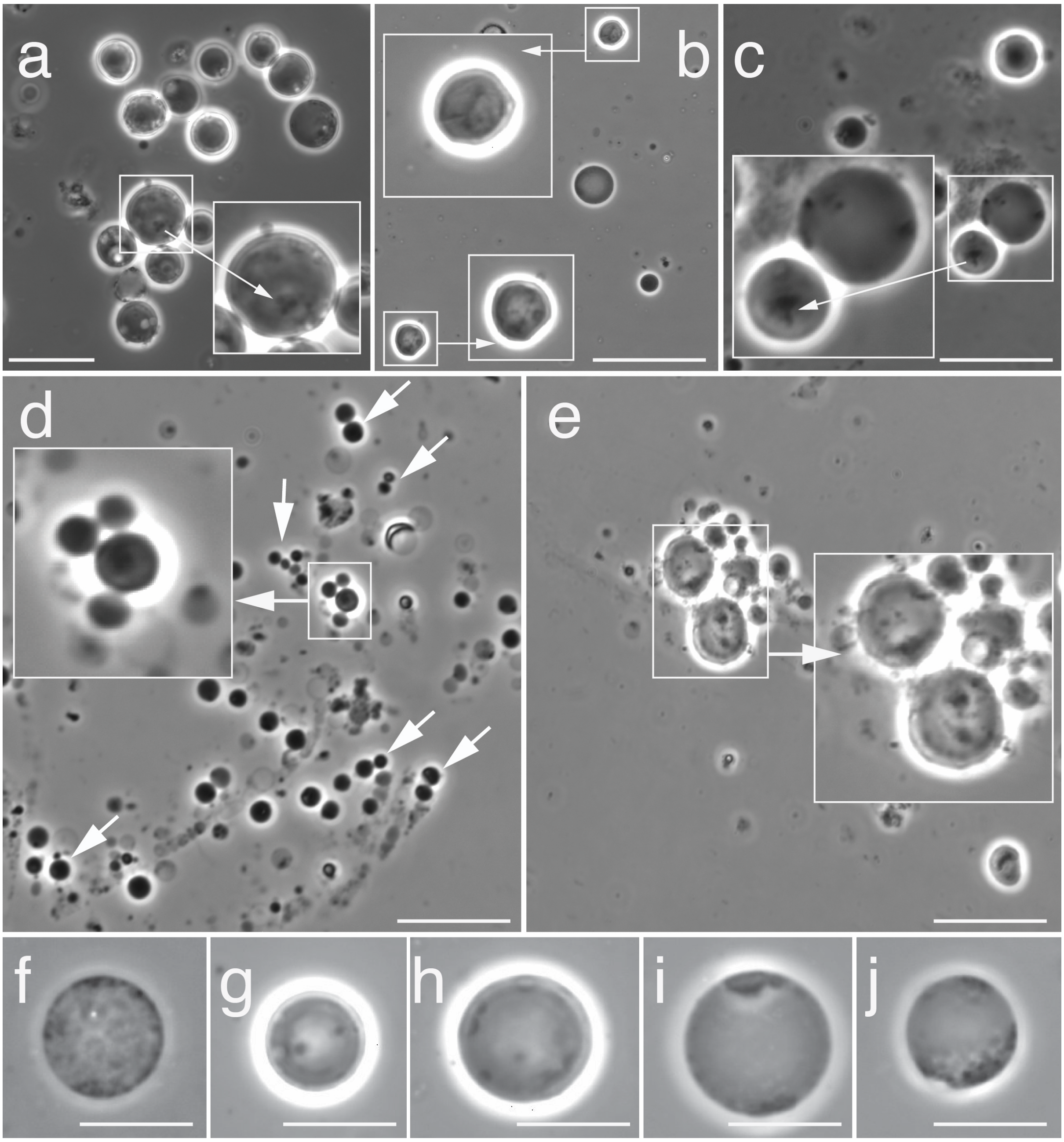


**Fig. S5 Log-phase *EM-P* cells with intracellular lipid droplets**

Images a-j show phase-contrast images of log-phase *EM-P* cells reproducing by binary fission (arrows) with intracellular accumulation of lipids. Given the smaller sizes of cells, such intracellular structures are barely visible in these cells. Highlighted regions in image a-e show magnified images of individual cells. Fig. S6 shows similar structures in larger *EM-P* cells kept in a continuous log phase. Scale bars: 5µm.

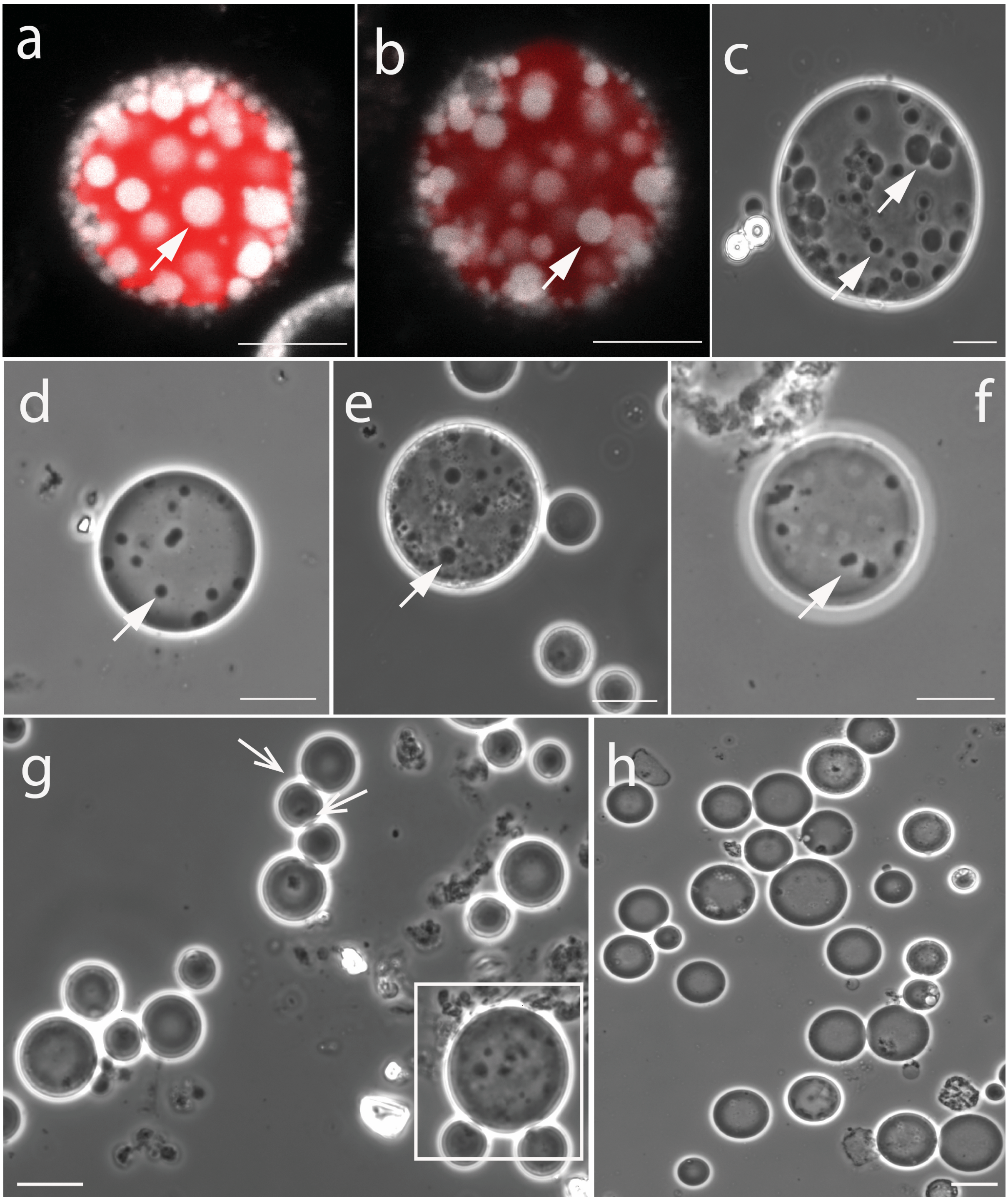


**Fig. S6** ***EM-P*** **cells kept in continuous log-phase**

Images a-f show STED and phase-contrast images of log-phase *EM-P* cells. Cells in these images were stained with universal membrane stain, FM$^{TM}$5-95 (white), and DNA stain, PicoGreen$^{TM}$ (red). Excess lipids accumulated as intracellular lipid droplets can be seen in images a & b (closed arrows). Open arrows in image-g points to cells reproducing by binary fission. The average sizes of these cells were considerably larger than the *EM-P* cells shown in Fig. 1, 2 & S4. Closed arrows in images a-f (highlighted region in image-g) point to lipid globules within *EM-P* cells. Open arrows in image-g point to cells reproducing by binary fission. Gradual transfer of lipid from droplets into the cell or vesicle membrane is shown in Fig. S7. Scale bars: 10μm.

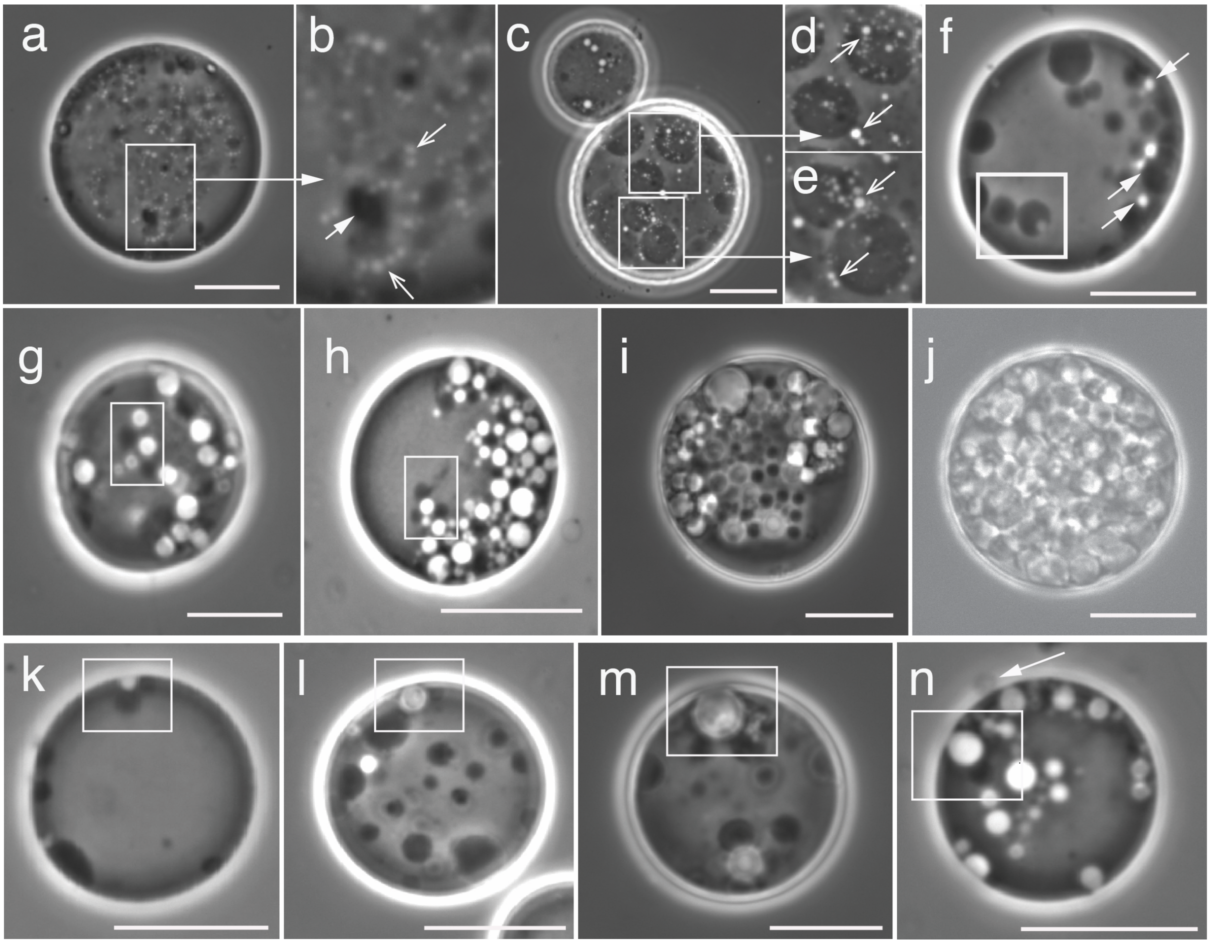


**Fig. S7 Transformation of lipid droplets into intracellular vesicles**

Images a-n show different stages involved in transforming lipid droplets into membrane structures within *EM-P*. All cells in this figure were kept in a prolonged log phase. Image-a shows an EM-P cell with tiny intracellular lipid globules. Image-b is the magnified region of *EM-P*'s cytoplasm, showing the presence of lipid droplets (closed arrows) and the formation of hollow vesicles (open arrows) at the periphery of these droplets. The gradual transformation of these tiny intracellular vesicles into hollow intracellular vesicles can be seen in images c-j. Images k-n show a similar formation of hollow vesicles closely associated with the cell membrane (highlighted regions). Arrow in image-n points to a hollow extracellular vesicle. Similar hollow extracellular vesicles or filamentous structures are also shown in Fig. 3. Scale bars: 10µm.

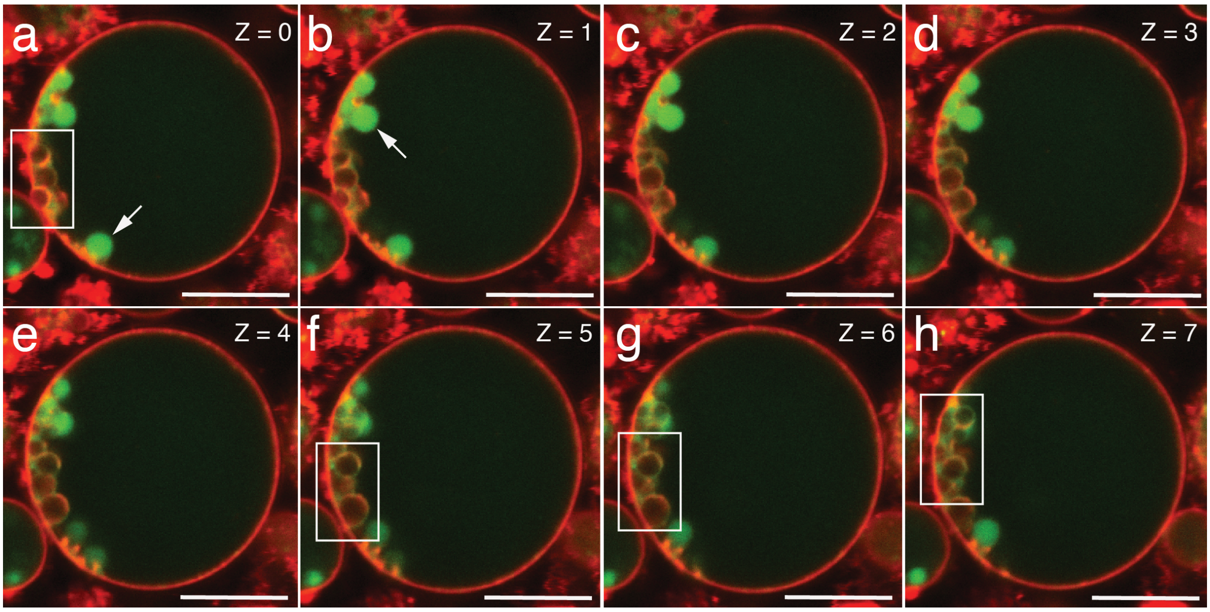


**Fig. S8 Transformation of $L_d$ membrane into intracellular vesicles**

Series of optical sections through late-log phase *EM-P* cells (0.2µm depth intervals). The cells were stained with universal membrane stain, FM™5-95 (all membrane, red), and *FAST*™ DiI ($L_d$-membrane specific, green). Scale bar: 10µm.

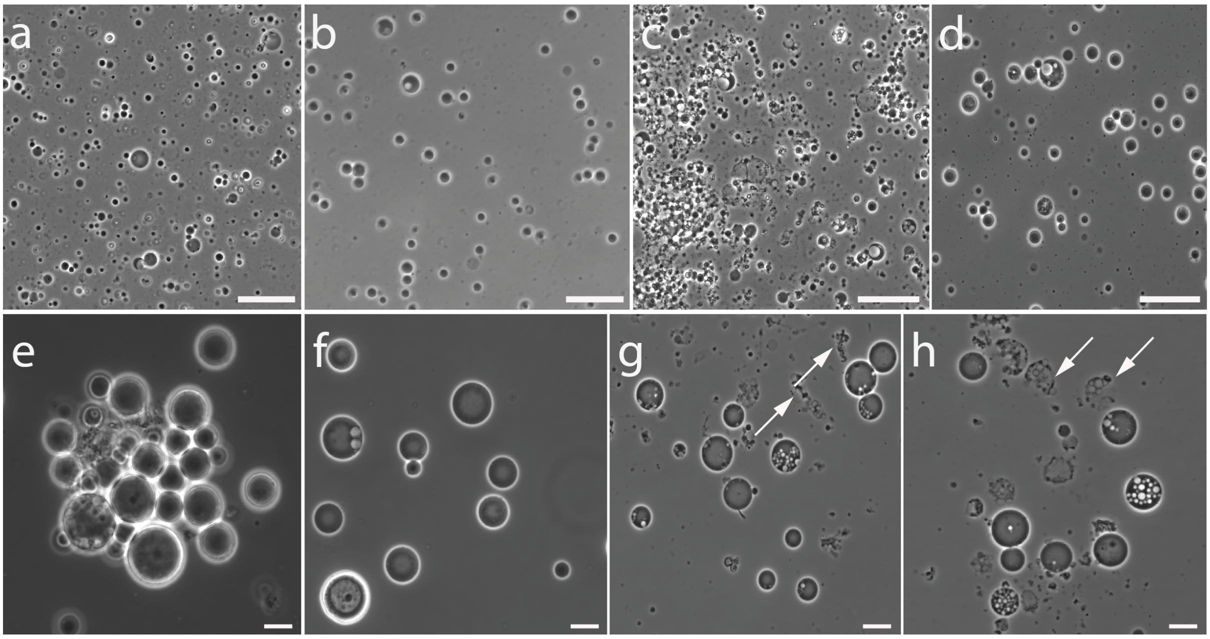


**Fig. S9 Influence of sudden decrease in media osmolarity on *EM-P*'s morphology**

Images a-d show late log-phase *EM-P* cells in 7%DSS-TSB. Images e-h show *EM-P* cells that were transferred from 7%DSS-TSB to 4%DSS-TSB. In comparison to the control incubations (a-d), the sizes of the cells in 4%DSS-TSB were significantly larger. We also noticed lysis of cells after the transfer, possibly due to the sudden changes in the osmolarity. Arrows in images g & h, points to the cells that underwent lysis. Scale bar: 10μm.

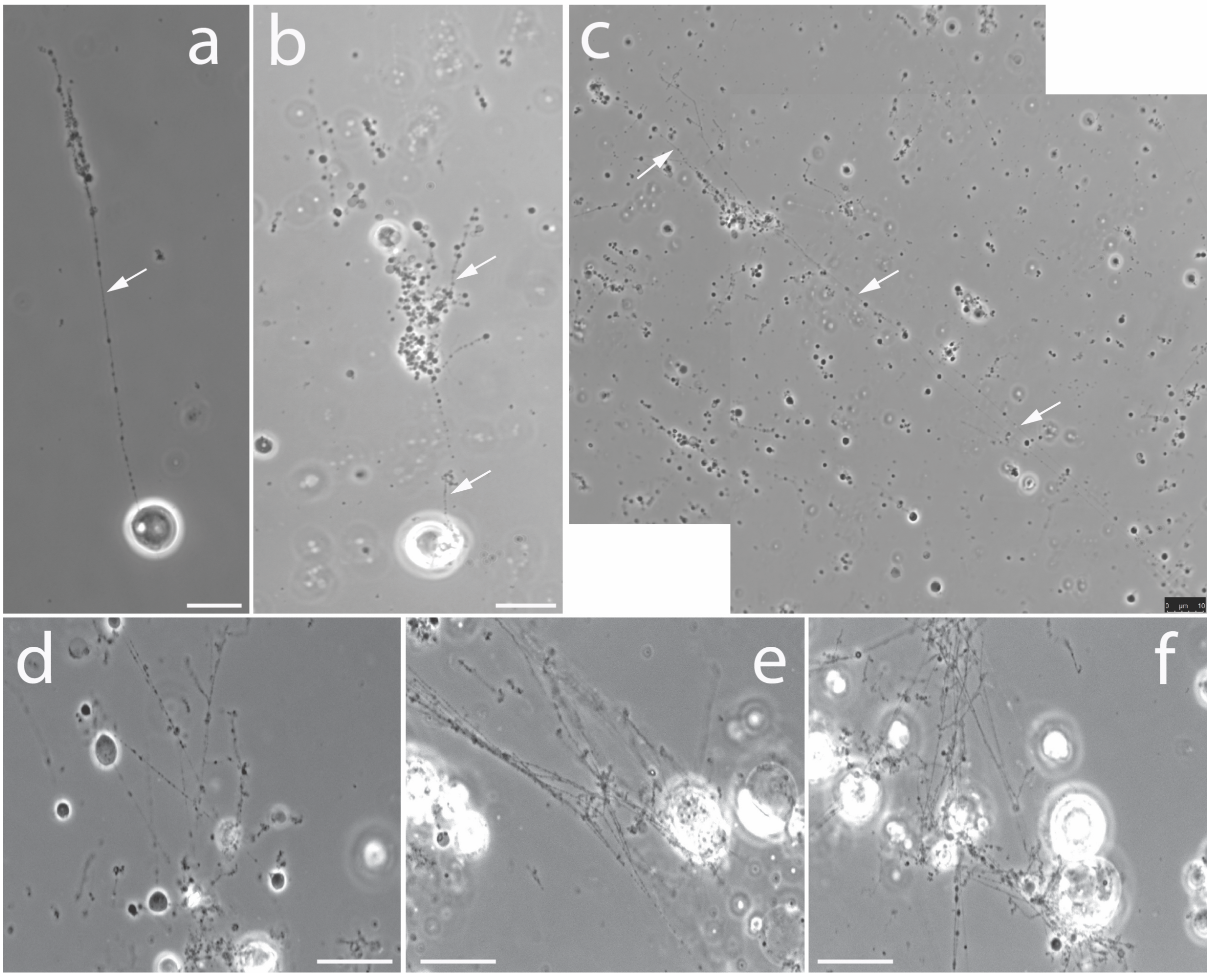


**Fig. S10 Influence of sudden increase in media osmolarity on *EM-P*'s morphology**

Images a-f show *EM-P* cells that were transferred from 7%DSS-TSB to 15%DSS-TSB. In comparison to the control incubations (Fig. S9 a-d), the sizes of the cells in 15%DSS-TSB were significantly smaller and contain longer filamentous extensions. Rather than being hollow, most filamentous extensions seems to have cytoplasm. Scale bar: 10μm.

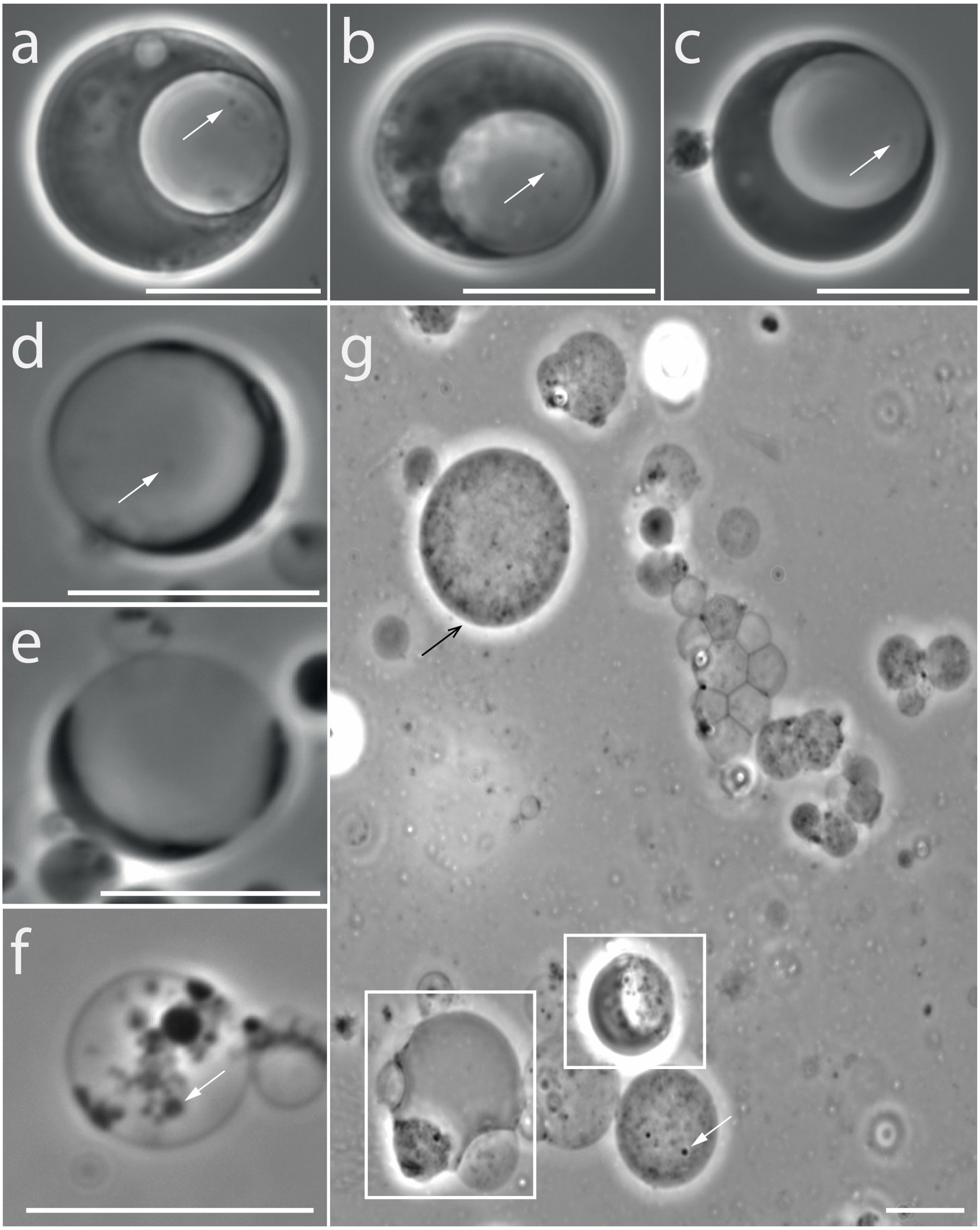


**Fig. S11 Morphological transformation of *EM-P* reproducing via the formation of internal daughter cells**

Images a-g show phase-contrast images of *EM-P* reproducing by formation of internal daughter cells. Image a-c show *EM-P* cells with intracellular vesicles. White arrows in these images highlight barely visible daughter cells (more clearly visible in images f, g, video 14 and 17). Images b-e show gradual depletion in the cytoplasmic content of the cell due to loss of cytoplasm to daughter cells. Image-f shows an *EM-P* cell completely depleted of its cytoplasmic content to daughter cells. Image-g shows *EM-P* cells in their last growth stage, when most of them transformed into spherical vesicles with tiny daughter cells (black arrow) and some cell with relatively little cytoplasm (Boxed cells). Lysis and release of these daughter cells is shown in Figure 6h, & Video 18. Scale bars: 10µm.

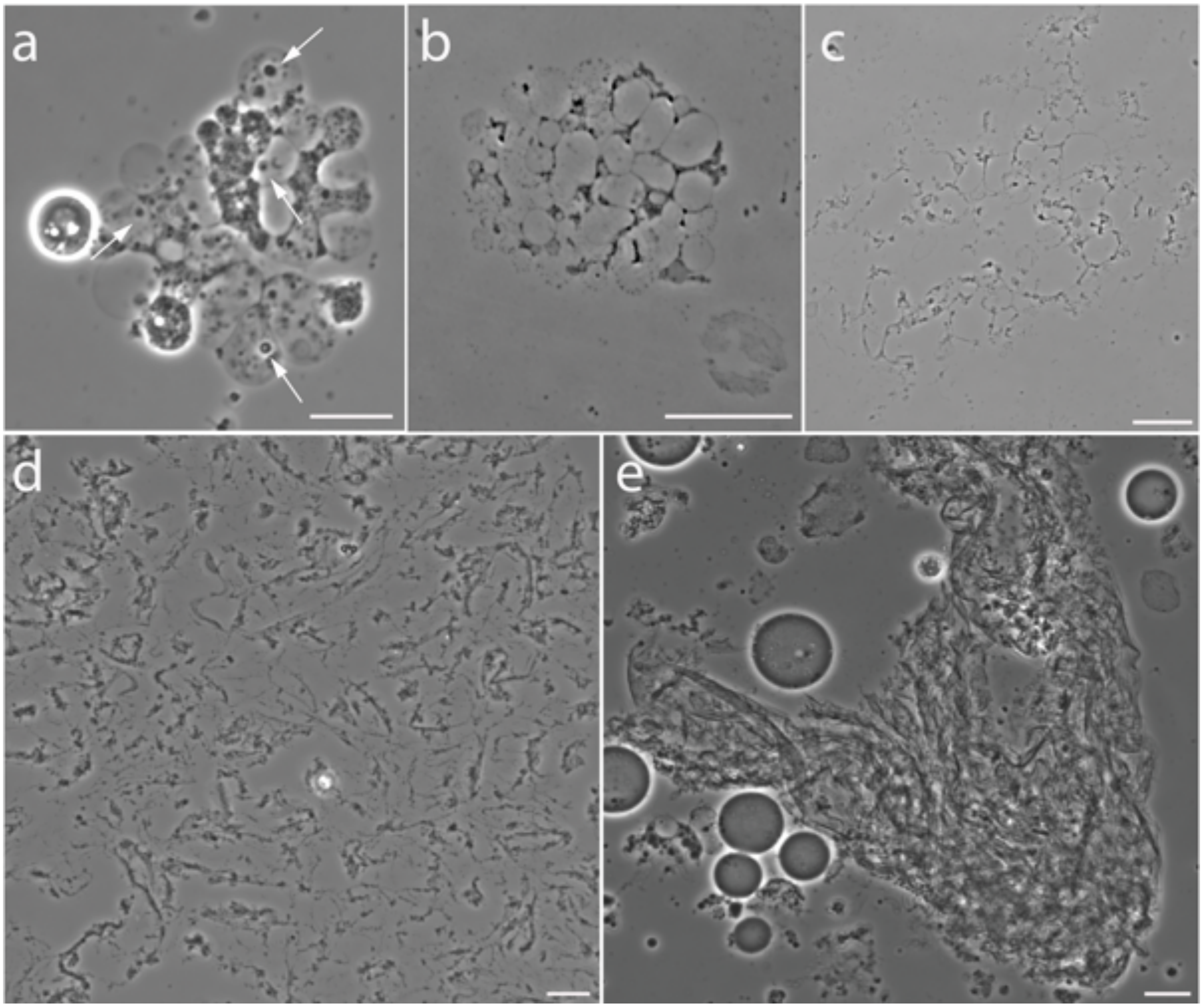


**Fig S12 Membrane debris observed during the stationary growth phase.**

Images a-e show sequential stages of membrane debris transformation. Image-a show intracellular vesicles released by cell lysis of *EM-P* cells (Video 15-18). Arrows in image-a point to tiny spherical daughter cells within these vesicles. Image-b shows vesicles that underwent lysis to release daughter cells (Movie S18). Image-c show the leftover membrane debris after the release of daughter cells. Images d & e show subsequent aggregation and transformation of membrane debris into fabric-like structures. Scale bars: 10μm.

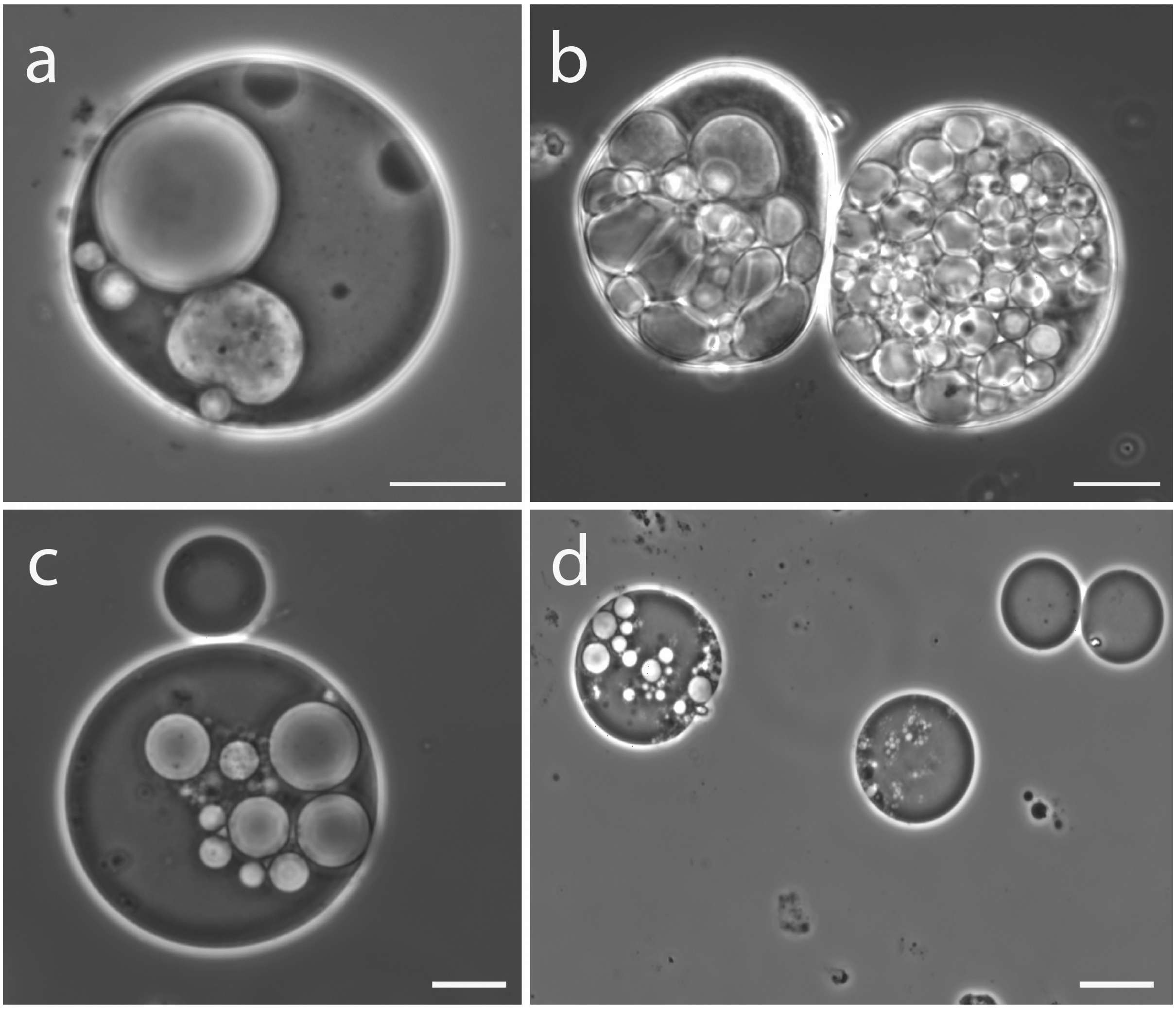


**Fig. S13** ***EM-P*** **cells reproducing via external daughter cells in 7%KCl-TSB**

Images a-d show the sequential morphological transformation of *EM-P* when grown in the presence of 7% KCl-TSB. Scale bars: 10µm.

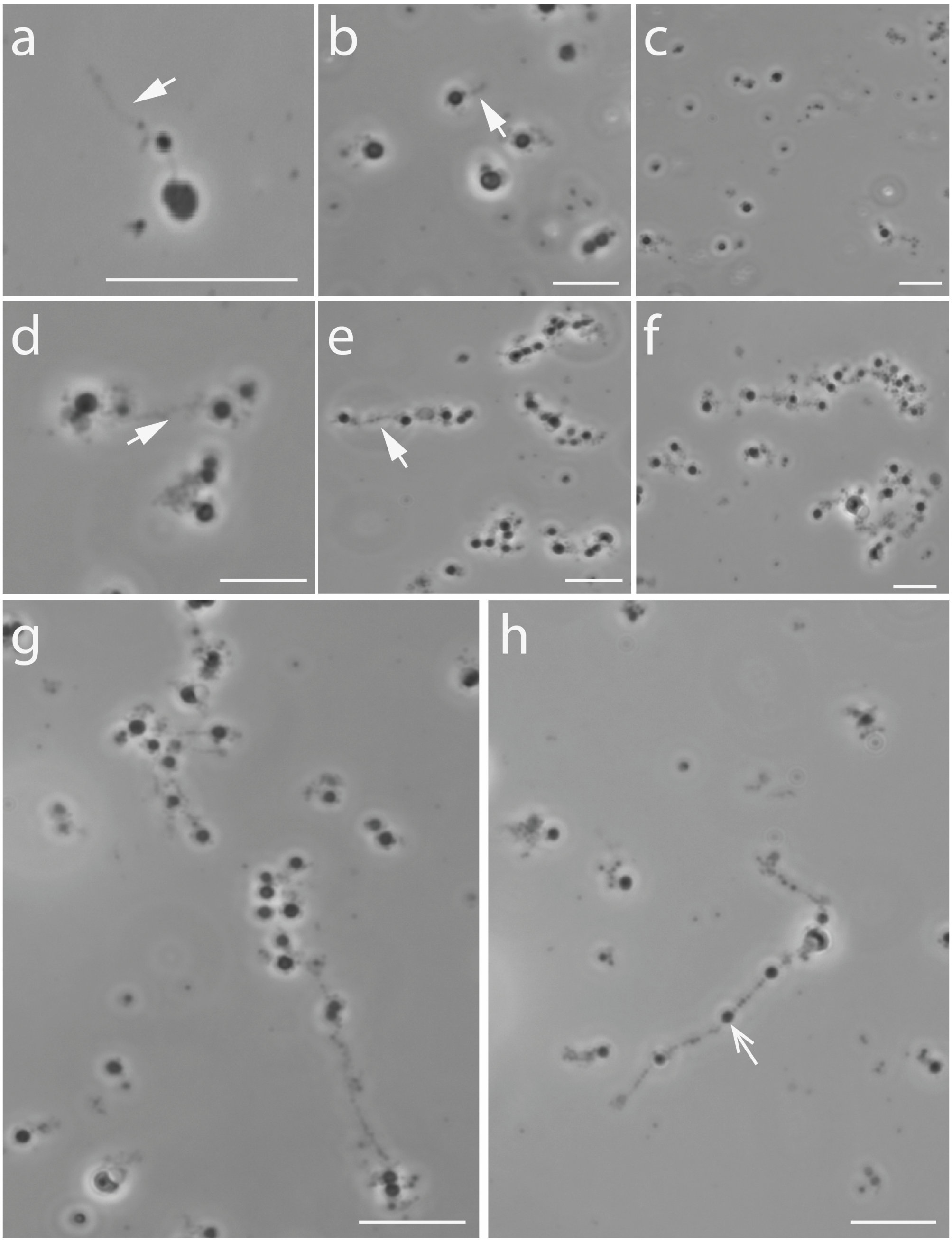


**Fig. S14 *EM-P* cells reproducing via external daughter cells in TSB-5%$MgCl_2$**

Images a-h show sequential morphological transformation of *EM-P* when grown in the presence of 5% $MgCl_2$-TSB. Closed arrows points to membrane tethers connecting individual daughter cells. Open arrows in these images' points to spherical daughter cells within the filaments (videos 20 & 21). Scale bars: 10µm.

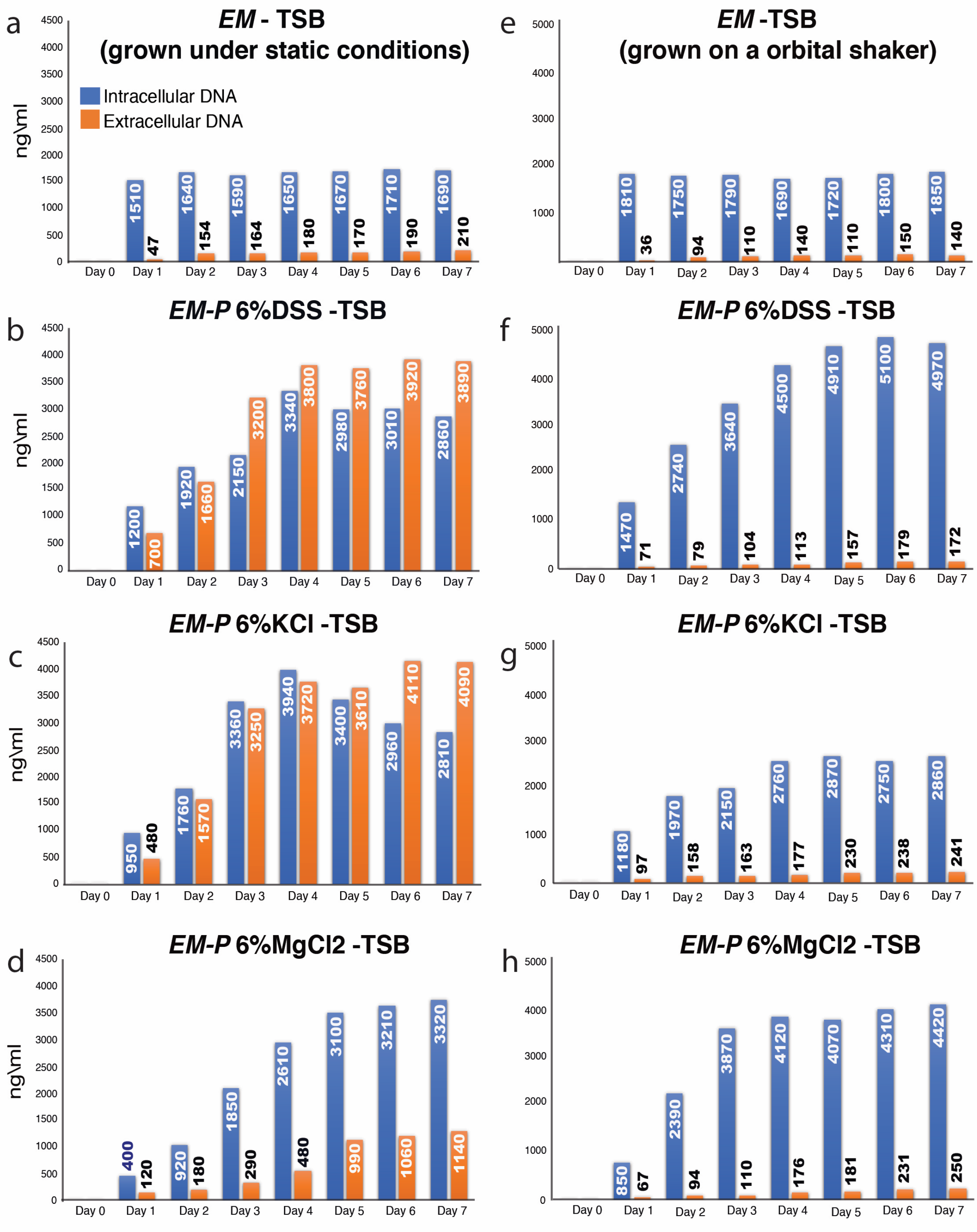


**Fig. S15 Reproductive efficiency of *EM-P***

Plots a-h show leakage of intracellular DNA during *EM-P*'s reproduction under different conditions. The plots (a-d) are the incubations done under static conditions. The salts composition of the media is shown above the individual graphs. Plot-a shows the results of wild-type *EM* (*Exiguobacterium strain M* with a cell wall). The rest of the plots b-d show *EM-P* grown in media containing DSS, KCl, and $MgCl_2$. The plots (e-h) are the incubations done under static conditions. Plot-e shows the results of wild-type *EM* (*Exiguobacterium strain M* with a cell wall) grown under static conditions. The rest of the plots b-d show *EM-P* grown in media containing DSS, KCl, and $MgCl_2$. All experiments are biological repetitions from different batches of *EM-P*'s inoculum (n=5).

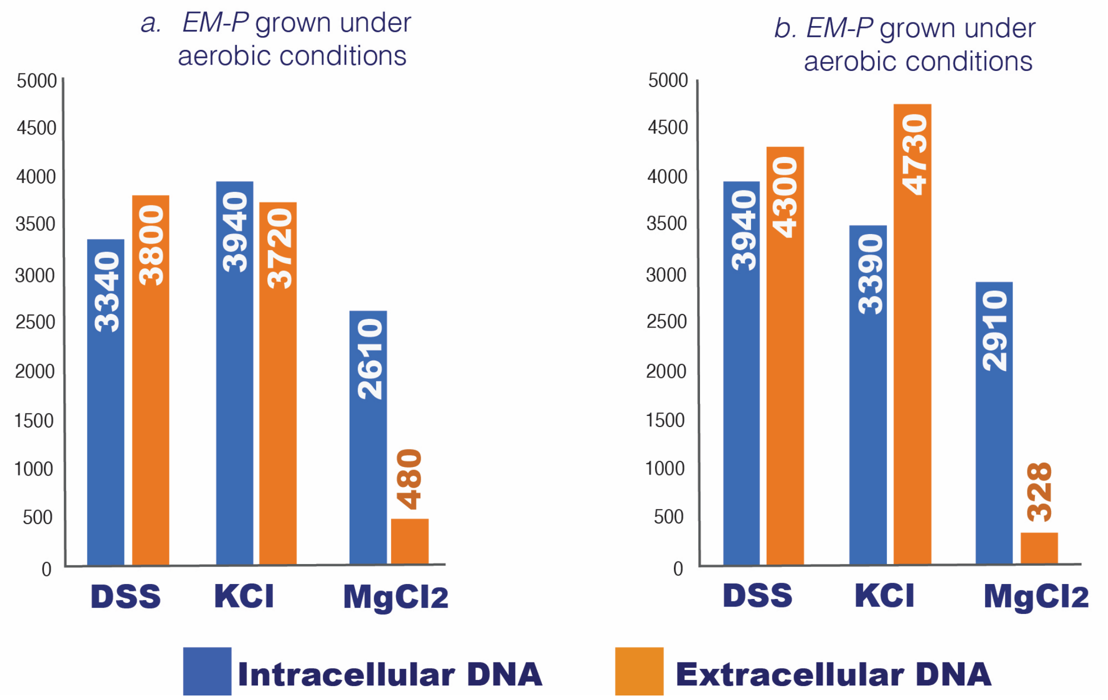


**Fig. S16 Reproductive efficiency of *EM-P* under aerobic and anaerobic conditions**

Plots a & b show leakage of intracellular DNA during *EM-P*'s reproduction under different media compositions under aerobic (a) and anaerobic conditions (b). Both experiments are conducted under static conditions. The salts composition of the media is shown above the individual graphs. All experiments are biological repetitions from different batches of *EM-P*'s inoculum (n=5).

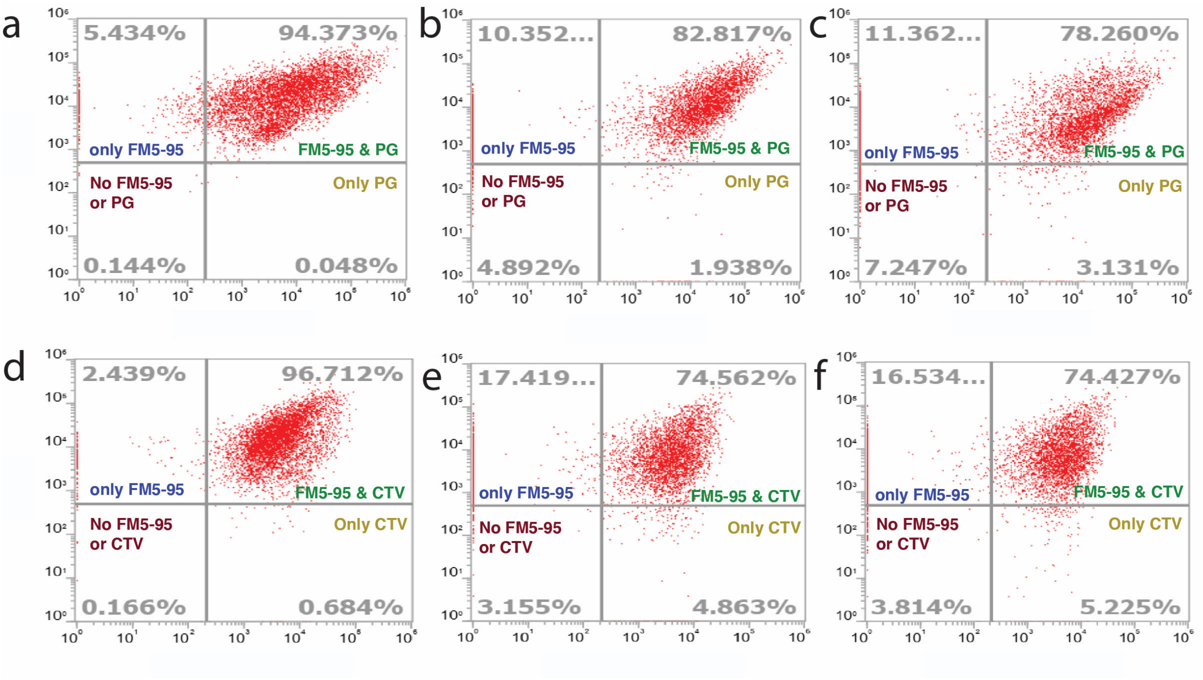


**Fig. S17 Viability of daughter *EM-P* daughter cells**

Figures a-f show flow cytometry quantification of different *EM-P* daughter cells. Cells from the stationary growth phase were passed through a 0.22µm filter to separate them from large parent cells and membrane debris (Fig. 7A). Subsequently, one set of these cells were stained with FM5-95 (FM5, membrane) and Pico Green (PG, DNA), to determine if daughter cells had an intact membrane and received genetic material from the parent cell. The second set of cells were stained with membrane dye FM5-95 (FM5, membrane) and Cell Trace Violet (CTV, intracellular esterase) to determine if the daughter cells had an intact membrane and exhibited cytoplasmic activity. A large fraction of cells in the top-right quadrant (FM5 & PG and FM5 & CTV) suggests most daughter cells are alive. Confocal images of these cells are shown in Fig. S18.

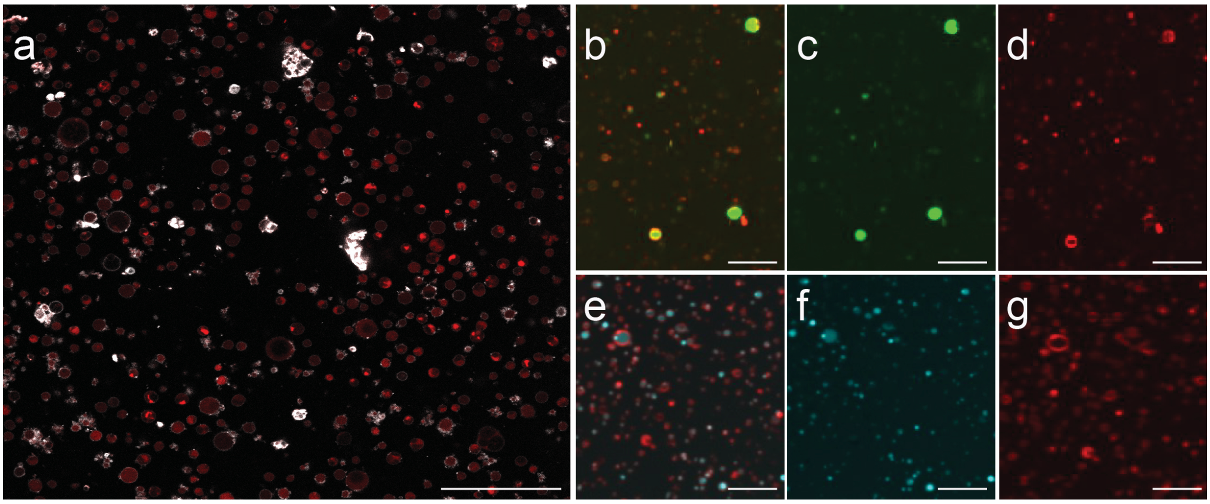


**Fig. S18** ***EM-P*** **cells in late stationery before and after filtration.**

Image-a shows a confocal microscope image of *EM-P* during the late growth stage. Cells in this image are stained with FM™5-95 (membrane, white) and PicoGreen™ (DNA, red). Images b & e show *EM-P* daughter cells after passing through a 0.45µm filter to separate them from larger parent cells and membrane debris. Cells in images b-d are stained with FM5™-95 (cell membrane, red) and PicoGreen™ (DNA, green). Images c & d show the same field of view in different channels specific for different dyes. Images e-g are images of *EM-P* daughter cells stained with FM5™-95 (cell membrane-red) and Cell Trace Violet™ (cytoplasm, cyan). Images f & g show the same field of view in different channels specific for different dyes. Scale bar: 10 µm (a) and 2µm (b-g).

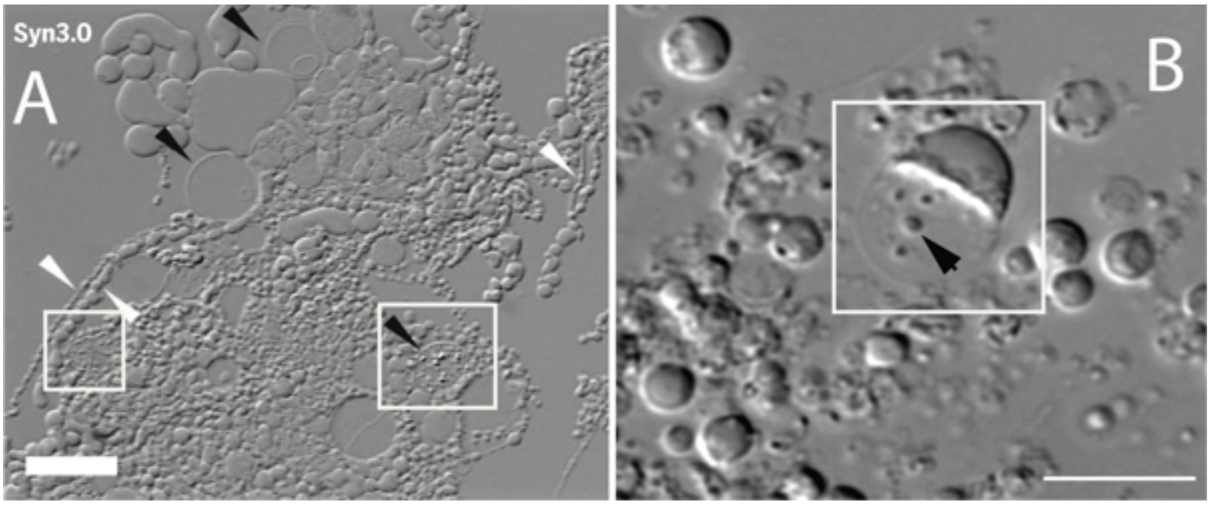


**Fig. S19 Morphological comparison of Syn 3.0 with *EM-P.*** Image-a: light microscope image of Syn 3.0 (originally published by Hutchison *et al.,* 2016, reproduced here with permission from AAAS) (72). Boxed regions in the image highlight hollow spherical cells with tiny internal globular structures. Image-b show morphologically similar *EM-P* cell. Highlighted regions in both the images show cells (both in Syn 3.0 and *EM-P*) with to tiny spherical inclusions (black arrows in image-b). Scale bars: 10µm (a & b).

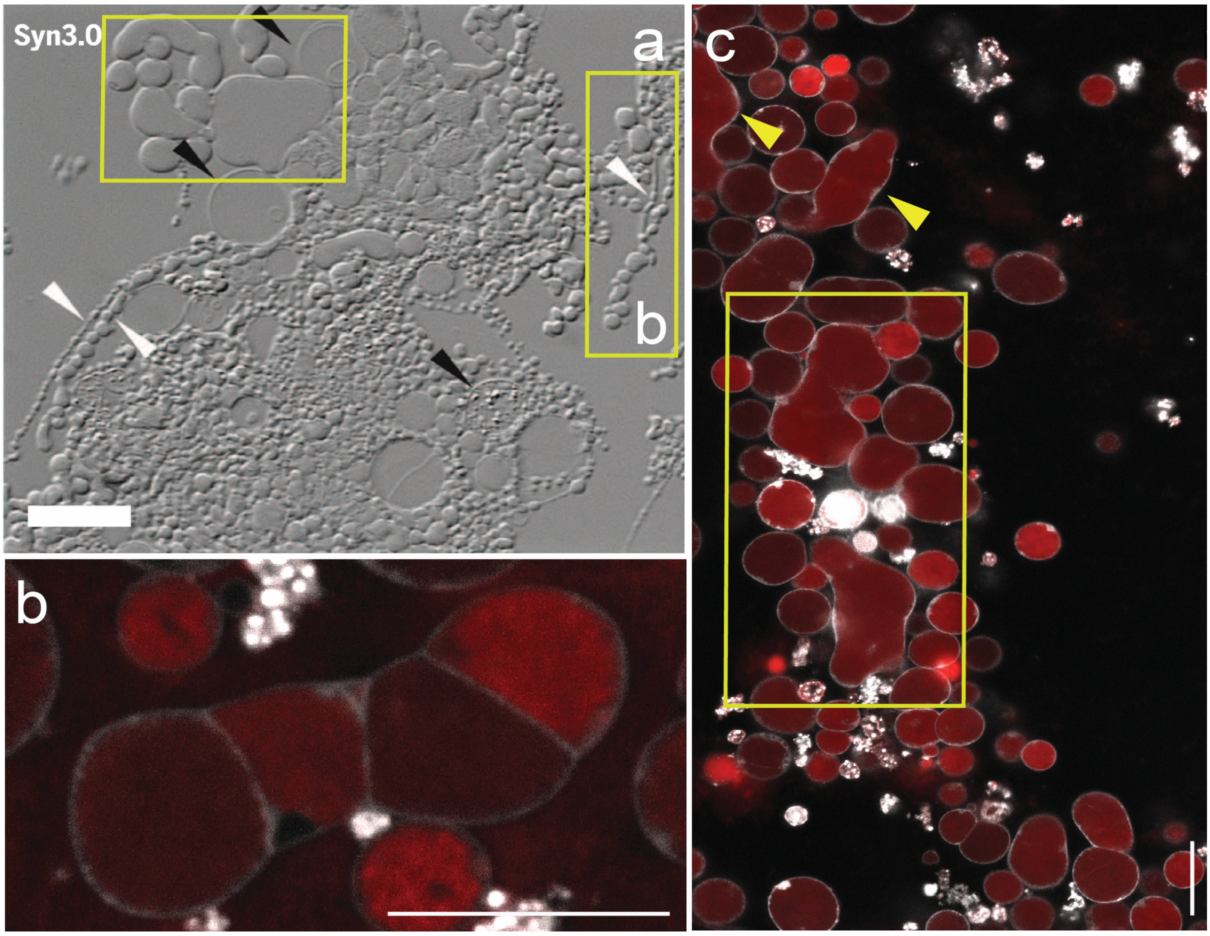


**Fig S20. Morphological comparison of Syn 3.0 with *EM-P*.**

Image-a: light microscope image of Syn 3.0 (originally published by Hutchison *et al.,* 2016, reproduced here with permission from AAAS) (72). Images b & c: STED microscope images of *EM-P*. Cells in these images are stained with FM™5-95 (membrane, white) and PicoGreen™ (DNA, red). Boxed regions (yellow) in image-a show Syn 3.0 that are of random morphology. *EM-P* cells of similar morphology can be seen in image-c. Image-b, shows an *EM-P* cells similar to the highlighted region in image-a (b). Scale bars: 10µm (a, & c) and 5µm (b).

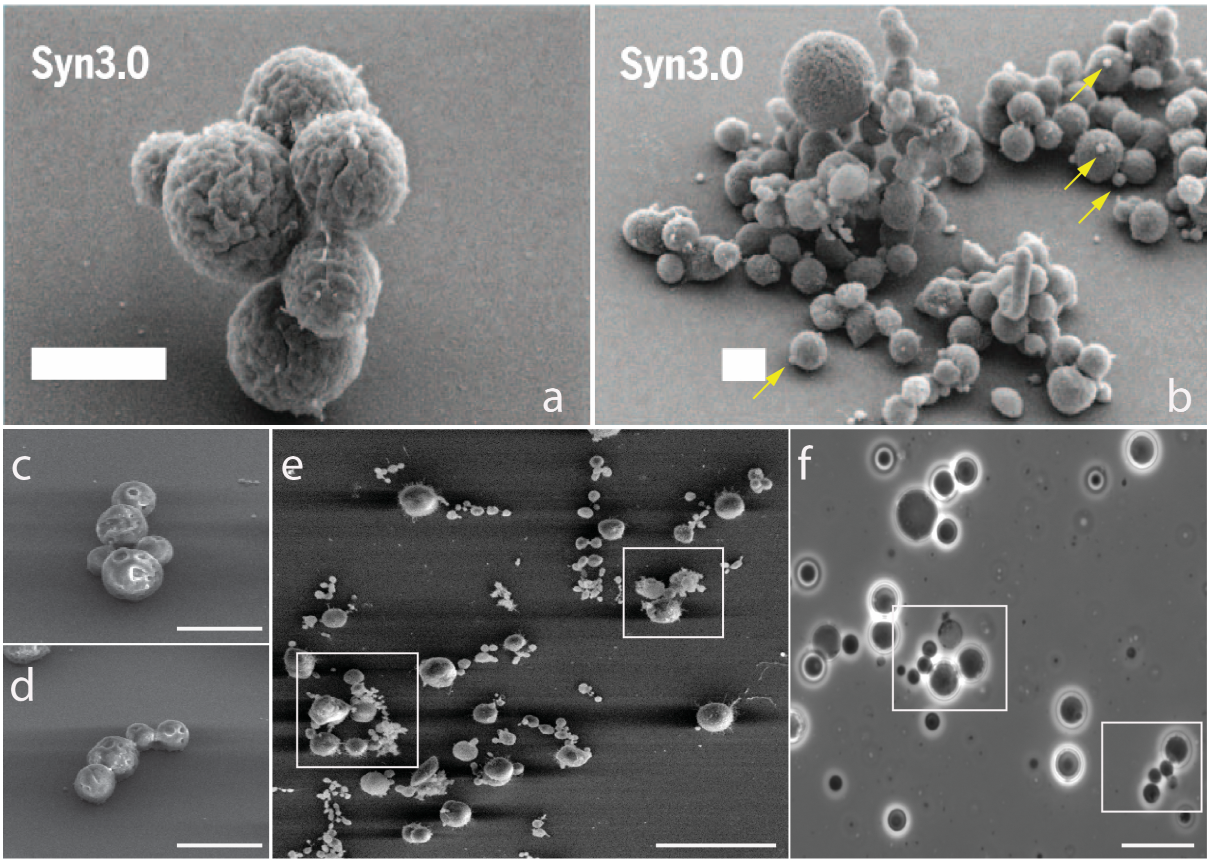


**Fig. S21 Morphological comparison of Syn 3.0 with *EM-P***

Images a & b are SEM images of syn3.0 (originally published by Hutchison *et al.,* 2016, reproduced here with permission from AAAS) (72). Images c-f are SEM and phase-contrast images of *EM-P*. Images c, d, and boxed regions in images e & f show clusters of *EM-P* cells of different sizes that were similar in their morphology to Syn 3.0 (a & b). Wrinkled surfaces of syn 3.0 suggest a state of excess membrane similar to *EM-P*. Yellow arrows in image-b point to tiny bud-like structures attached to Syn 3.0. A similar mechanism of reproduction by budding was also observed in *EM-P* and shown in Fig. 1 & 2. Scale bars: 1µm (a & b), 5µm (c-e) and 10µm (f).

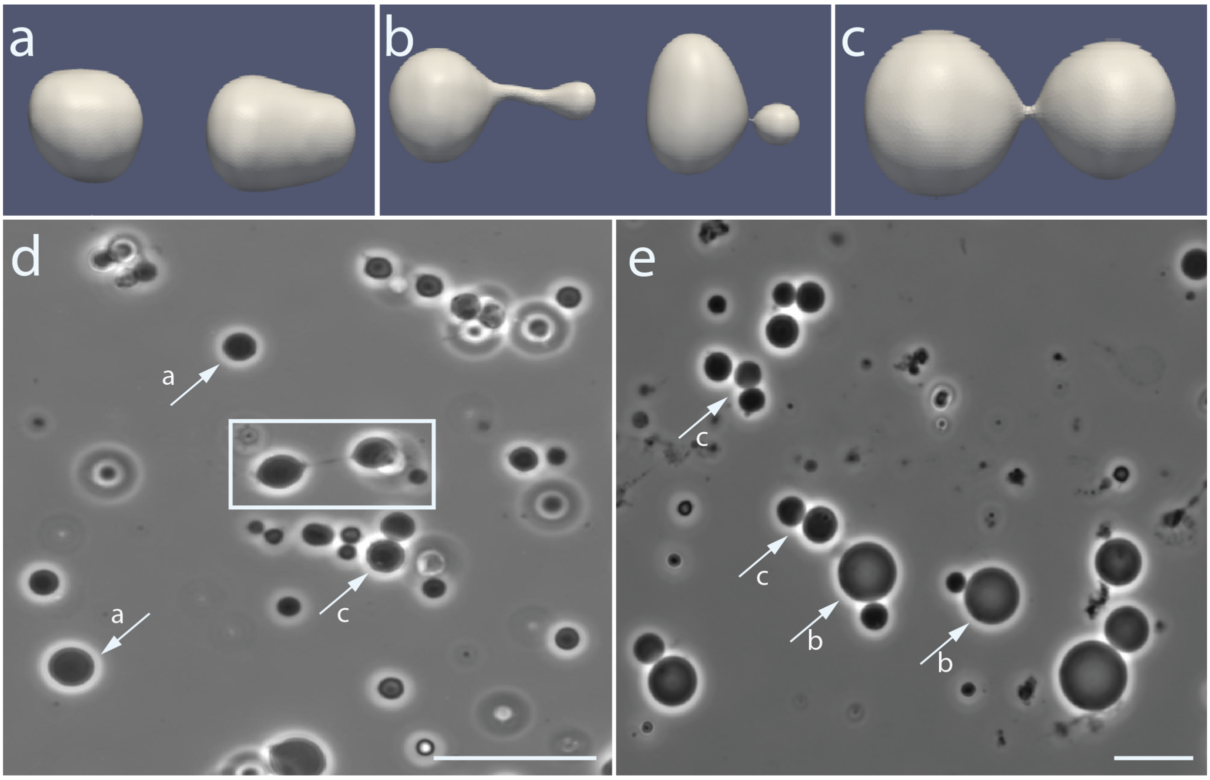


**Fig. S22 Comparison of *EM-P* morphology with theoretical protocell morphologies**

Images a-c show theoretically predicted morphologies of protocells (originally published by Ruiz-Herrero et al., 2019) (73). Reproduced here with permission from the American Physical Society. Images d & e show morphologically analogous *EM-P* cells. Images a-c show spherical cells undergoing symmetric (c) or asymmetric (b) cell division. Arrows in images d & e point to morphologically similar cells. The boxed region in image-d shows *EM-P* cells that underwent binary fission but are still connected by a narrow membrane connection (video 3). *EM-P* cells undergoing a similar reproduction, either by budding or binary fission, can be seen in Fig. 1 & 2. Scale bars: 10μm (d & e).

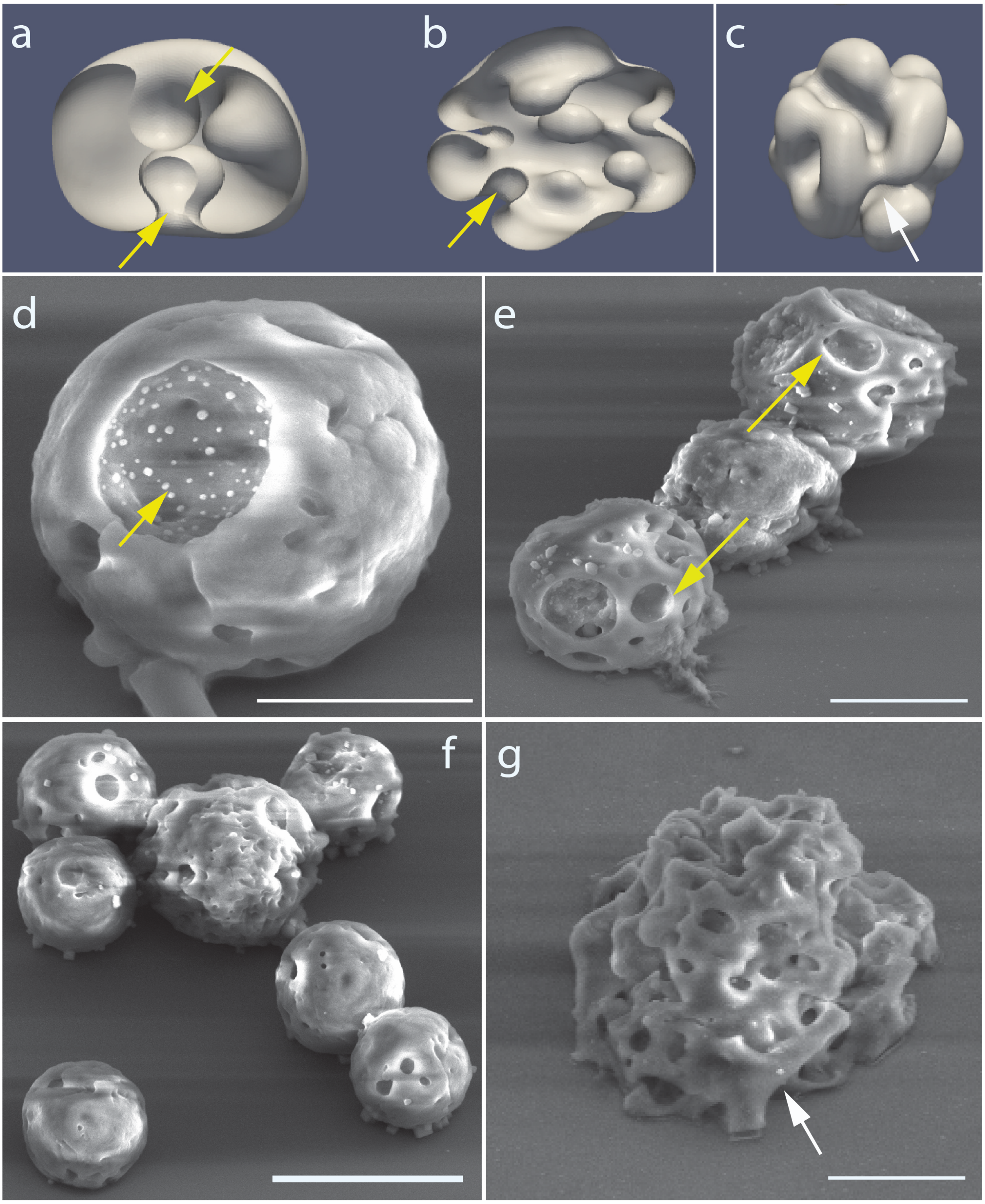


**Fig. S23 Comparison of *EM-P* morphology with theoretical protocell morphologies**

Images a-c show theoretically calculated (originally published by Ruiz-Herrero et al., 2019. Reproduced here with permission from the American Physical Society) (73). Images d-g are morphologically analogous to *EM-P* cells. Images a & b show cells forming surface depressions to accommodate excess membrane. Images d & e show similar surface depressions in *EM-P* (Yellow arrows). Image-f shows a cluster of *EM-P* cells with similar surface depressions, which eventually transformed into intracellular vesicles, as shown in Fig. 6. Image-g shows an *EM-P* cell with surface membrane folds, resembling the theoretically predicted morphologies (c). Arrows in images c & g point to theoretically predicted Y-shaped folding pattern. Scale bars: 2µm (d), and 5µm (e-g).
